# Age-related STAT3 signaling regulates severity of respiratory syncytial viral infection in human bronchial epithelial cells

**DOI:** 10.1101/2023.09.20.558606

**Authors:** Caiqi Zhao, Wei Wang, Yan Bai, Gaurang Amonkar, Hongmei Mou, Judith Olejnik, Adam J Hume, Elke Mühlberger, Yinshan Fang, Jianwen Que, Rachel Fearns, Xingbin Ai, Paul H. Lerou

## Abstract

Respiratory syncytial virus (RSV) can cause severe disease especially in infants; however, mechanisms of age-associated disease severity remain elusive. Here, employing human bronchial epithelium models generated from tracheal aspirate-derived basal stem cells of neonates and adults, we investigated whether age regulates RSV-epithelium interaction to determine disease severity. We show that following RSV infection, only neonatal epithelium model exhibited cytopathy and mucus hyperplasia, and neonatal epithelium had more robust viral spread and inflammatory responses than adult epithelium. Mechanistically, RSV-infected neonatal ciliated cells displayed age-related impairment of STAT3 activation, rendering susceptibility to apoptosis, which facilitated viral spread. In contrast, SARS-CoV-2 infection of ciliated cells had no effect on STAT3 activation and was not affected by age. Taken together, our findings identify an age-related and RSV-specific interaction with neonatal bronchial epithelium that critically contributes to severity of infection, and STAT3 activation offers a potential strategy to battle severe RSV disease in infants.

## INTRODUCTION

Infection of human bronchial epithelium by respiratory syncytial virus (RSV) occurs in people of all ages; however, only certain populations, including infants younger than 6 months of age, are at risk of developing severe RSV disease. RSV infection is the leading cause of acute and severe lower airway disease in young children and the primary cause of hospitalization of infants in the first year of life worldwide^1,2^. The pathophysiology of infant RSV disease is characterized by epithelial cell apoptosis and sloughing, inflammation, mucus hyperplasia, and airway obstruction^3,4^. Beyond acute morbidity, RSV infection in infancy is a major risk factor for hyperreactive airway disease and asthma later in life^5–10^. To protect the very young from severe RSV disease, decades of intensive effort was spent on vaccine development for infants with disappointing results^11–14^. Very recently, a vaccine for administration during late pregnancy and an RSV antibody^15–18^ were clinically approved, offering promising prophylactic strategies against RSV in newborns and infants. However, the challenge of identifying effective treatment strategies for active RSV infection in infants remains.

Most studies of age-related RSV disease focus on the immature immune system in infants^3,19^. Apart from immune cells, human bronchial epithelium is the first contact of RSV^20–22^ and plays a central role in orchestrating the inflammatory response following respiratory viral infection. The age of the host may regulate the interaction between RSV and bronchial epithelial cells and thus contribute to age-related severity of RSV disease. This possibility is accentuated by the contrasting predilection for severe disease with age following infection by SARS-CoV-2. Like RSV, SARS-CoV-2 also targets ciliated cells^23,24^. However, in contrast to RSV infection that mostly affects infants, severe COVID-19 disproportionally affects adults, and infants and young children typically have no symptoms after SARS-CoV-2 infection making them possible spreaders of ongoing and emerging virus variants^25^. The drastic difference in clinical outcomes of infants and adults following infection by RSV and SARS-CoV-2 supports the hypothesis that there are virus-specific interactions with human bronchial epithelium that differ between age groups and influence disease outcome.

The mechanisms that mediate age-related susceptibility to severe RSV disease remain elusive. This is attributable to limitations in pre-existing experimental models. A narrow species tropism of RSV creates a significant obstacle in modeling infant RSV disease using animal models^26^. In addition, the *in vitro* models, such as immortalized human HEp-2 and A549 cell lines, poorly represent human airway epithelium that is polarized and contains multiple cell types. To capture the structural and cellular complexity of human bronchial epithelium *in vivo,* air-liquid interface (ALI) culture of human bronchial basal stem cells (BSCs) is widely used^27,28^. Human bronchial BSCs are traditionally isolated from lung biopsy and airway brushing samples that involve invasive procedures and are often contraindicated in infants. For this reason, prior ALI models of RSV infection utilized bronchial BSCs from children and adults^8,20,29,30^. These previous studies showed that RSV infection has no cytopathic effect on adult epithelial cells in ALI but causes cell sloughing and apoptosis in pediatric ALI cultures^20,29,30^. To address RSV infection in infants, ALI cultures of nasopharyngeal BSCs, which are more accessible than bronchial BSCs, have been employed^30,31^. However, nasopharyngeal BSCs differentiate into upper respiratory epithelial cells that display lower cytopathic and inflammatory responses to RSV compared to bronchial epithelial cells^30^. These findings indicate the limitation of the utility of nasopharyngeal BSCs in modeling RSV infection. Thus, while previous findings provide evidence for age-related epithelial infection by RSV, specifically how infant bronchial epithelium responds to RSV, and mechanisms determining age-related differences remain unknown.

To generate a clinically relevant human bronchium model for the investigation of age-related severe RSV disease in infants, we employed an established methodology of BSC derivation from tracheal aspirate (TA) samples^32–35^. TA samples can be collected as part of routine care for intubated patients and are typically considered medical waste. TA contains rare BSCs that can be expanded in culture and differentiated into multiple bronchial epithelial cell types^32,36^. Importantly, TA BSCs are indistinguishable from bronchial biopsy BSCs in transcriptomic profiles and differentiation potentials^33,35,36^ and can reproduce bronchial epithelium phenotypes found *in vivo*, shown by our previous studies of congenital diaphragmatic hernia in newborns^34^ and COVID-19 in adults^35^. Here, employing bronchial epithelium models generated from neonatal and adult TA BSCs, we investigated the relationship between age and epithelial cell response to infection by RSV. We also tested SARS-CoV-2 infection as a potential contrast for age-related differences in epithelial response. Our data indicate that insufficient STAT3 signaling in neonatal ciliated cells following RSV infection, compared to the adult counterpart, mediates significant cytopathy and severe infection. In addition, STAT3 regulation of age-related severity of human bronchial epithelium infection is specific to RSV.

## RESULTS

### Human bronchial epithelium models generated from TA BSCs of neonates and adults reproduce age-related severity of RSV infection

To circumvent technical difficulties of BSC isolation from infants younger than 6 months of age, we derived BSCs from TA samples of full-term newborns who were intubated for primarily non-pulmonary diseases (Supplementary Table 1) using an established Rho/Smad/mTOR inhibition culture condition^32,33^. Neonatal TA BSCs differentiate into functional epithelial cells in ALI for 3 weeks (D21) (Figures 1A and 1B)^32,33^, a time point when epithelial differentiation of BSCs is considered complete^37,38^. As controls, TA BSC lines from adult patients (32-55 years of age) who were intubated due to cardiogenic and neurogenic respiratory failure were similarly derived (Supplementary Table 1) and differentiated in ALI (Figures 1A and 1B). Transcriptome profiling of neonatal and adult TA BSCs identified differentially expressed genes enriched in the cell cycle pathways (Figures S1A-S1C), consistent with differences in BSC proliferation with age^39^. Neonatal and adult TA BSCs exhibited similar differentiation potentials in ALI, evidenced by comparable percentages of all major epithelial cell types between the two ages, including RFX3^+^ ciliated cell (41.4%±3.1% (neonate) vs 38.7%±1.7% (adult), p=0.53), SCGB1A1^+^ club cells (10.0%±1.9% (neonate) vs 15.1%±3.9% (adult), p=0.22), MUC5AC^+^ goblet cells (9.7%±2.5% (neonate) vs 10.3%±2.4% (adult), p=0.88), and the remaining TP63^+^ BSCs (26.1%±3.8% (neonate) vs 27.6%±3.5% (adult), p=0.79) (Figures 1B and S1D). Ciliated cells, which are the cell target of both RSV and SARS-CoV-2, constituted approximately 70% of apical surface cells in both neonatal and adult ALI cultures (Figure 1B). Antibody staining of human donor lungs also showed no difference in the abundance of ciliated cells in the intrapulmonary bronchi of infants and adults (Figures S1E and S1F). Taken together, our and previous findings^36,39^ show that age has no effect on cellular composition of human bronchial epithelium *in vivo* or differentiation potentials of BSCs *in vitro*.

**Figure 1.**
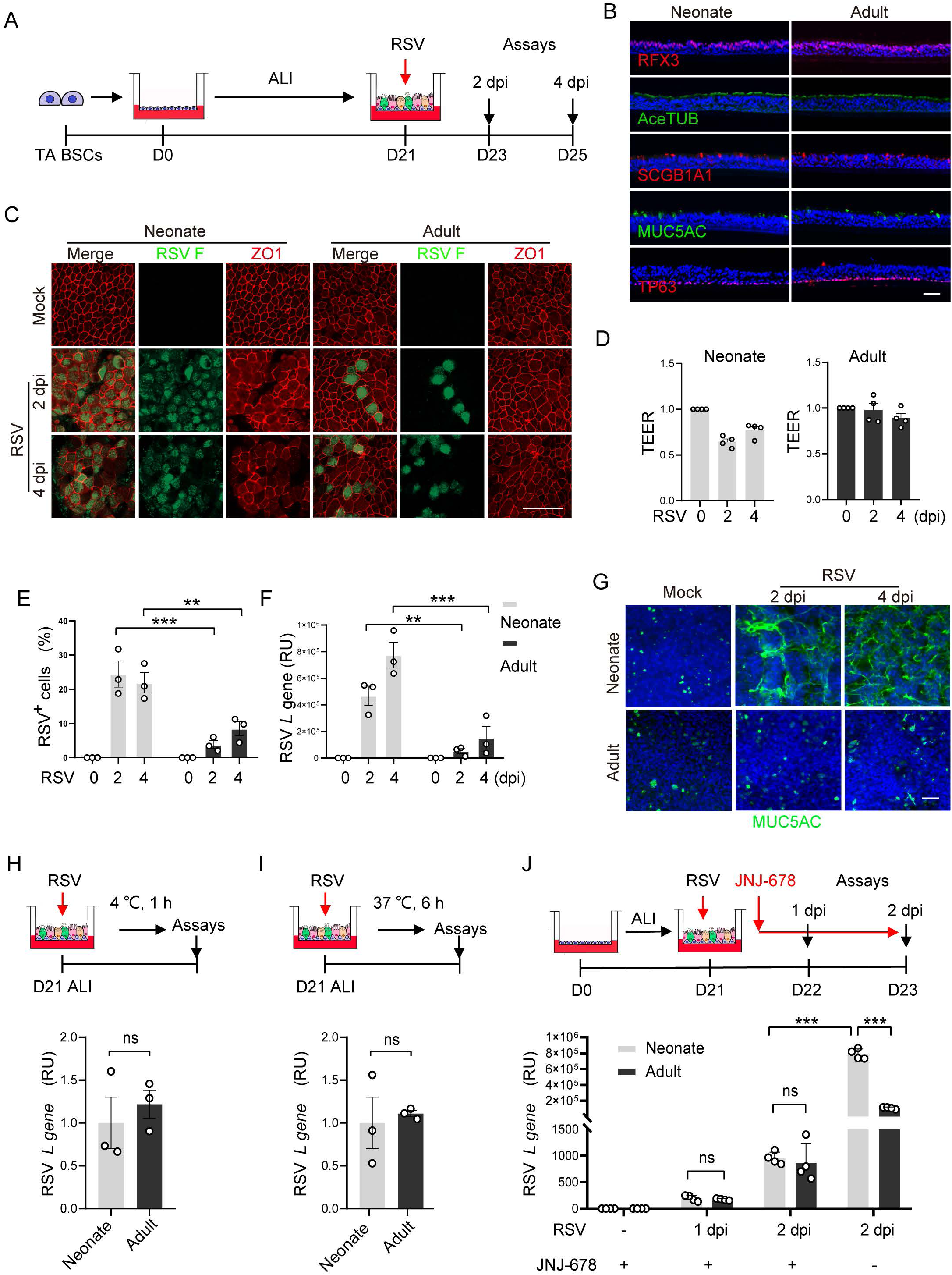
Human bronchial epithelium models generated from TA BSCs of neonates and adults show age-related RSV infection. (A) Schematic of RSV infection of day 21 (D21) ALI cultures of neonatal and adult TA BSCs in 24-well inserts (n=4 BSC lines for each age group). RSV strain A2 was applied apically (MOI 2, 4X10^5^ PFU) for 1 hour. Assays were performed at 2 and 4 dpi and presented in (B-G). (B) Representative double staining for markers of major epithelial cell types including RFX3 and AceTUB (acetylated tubulin) for ciliated cells, SCGB1A1 for Club cells, MUC5AC for goblet cells, and TP63 for remaining BSCs. (C) Representative whole-mount double staining for RSV F protein and ZO1. (D) TEER measurements normalized to baseline without RSV infection. (E) The relative abundance of RSV F^+^ cells in ALI cultures. (F) The relative level of RSV *L* gene by RT-qPCR. (G) Representative whole-mount staining for MUC5AC. (H) Assays of RSV binding at 4℃ for 1 hour followed by quantification of relative RSV *L* gene levels by RT-qPCR. (I) Assays of RSV early transcript and/or genome replication after infection at 37℃ for 6 hours by RT-qPCR. (J) Assays of JNJ-678 (100 nM) treatment at 6 hpi followed by quantification of relative RSV *L* gene levels by RT-qPCR at 1 and 2 dpi. Each dot represents one BSC line. Bar graphs represent mean ± SEM. Statistical significance was calculated by two-way ANOVA followed by Sidak’s multiple comparison test in (E, F, and J) and by Student’s t-test (two-tailed) in (H and I). **p<0.01, ***p<0.001, and ns, not significant. Scale bars, 50 μm.

To test whether RSV infection of human bronchial epithelium was affected by age, we infected day 21 ALI cultures of neonatal and adult TA BSCs with RSV strain A2. Prior to infection, the apical surface of ALI cultures was subjected to multiple saline washes to remove mucus. RSV (4X10^5^ PFU) was applied apically for 1 hour followed by saline washes to remove unbound viral particles. Viral load and epithelial responses were assayed at 2 days post-infection (dpi) and 4 dpi (Figure 1A). We found that RSV infection of adult ciliated cells elicited no cytotoxicity or barrier damage but induced ciliary dyskinesia (Figures 1C, 1D, S2A, and S2B), similar to previous reports^20,29,40,41^. In contrast, RSV infection caused cell sloughing and barrier damage in neonatal ALI cultures, manifested by loss of ZO-1, reduced transepithelial electrical resistance (TEER), and crater-like areas in the apical surface (Figures 1C, 1D, and S2C). Compared to adult cultures, neonatal ALI cultures also had more RSV-infected cells (>3-fold) and higher RSV RNA levels (>10-fold) (Figures 1E, 1F, and S2D). Furthermore, only neonatal ALI cultures showed mucus hyperplasia following RSV infection (Figures 1G, S3A, and S3B) that was associated with an increase in expression of inflammatory signals, including CXCL10, *TNFA*, and interferon-related genes (Figures S3C and S3D). Lastly, age-related severe infection of neonatal epithelial cells was also induced by a clinically relevant human RSV B strain (WV/14617/85) (Figure S4). Taken together, the neonatal bronchial epithelium model of RSV infection captures clinical hallmarks of severe RSV disease in infants^3,4^ and thus provides a viable system to investigate how age affects epithelial cell interaction with RSV.

### Human infant bronchial epithelial cells support RSV spread

To test whether RSV binding to bronchial epithelial cells is affected by age, we incubated RSV with neonatal and adult ALI cultures at 4℃ for 1 hour (Figure 1H), a condition that supports RSV binding but not viral entry into host cells. After washes to remove unbound viral particles, we performed viral RNA assays and showed that the amount of virus bound to epithelial cells was similar between neonatal and adult ALI cultures (Figure 1H). Therefore, age has no effect on RSV binding to human bronchial epithelial cells.

After viral entry into ciliated cells, RSV undergoes genome expression and replication to generate new virions. At 4–6 hours post infection (hpi), new viral RNA and protein can be detected intracellularly, but new RSV virions are not yet assembled or released until ∼12 hpi^42^. Leveraging this time course of RSV infection, we assayed intracellular viral RNA levels at 6 hpi and found no difference between neonatal and adult ALI cultures (Figure 1I). These findings indicate that age has no effect on primary expression and/or replication of RSV genome in the infected ciliated cells. As such, age-related differences in RSV infection of human bronchial epithelial cells likely occur at a later time in the infection cycle.

Next, we tested whether age affects the spread of newly released virions. To do so, we treated neonatal and adult ALI cultures with JNJ-678 which blocks RSV fusion^43^ and thereby prevents released virions from infecting other cells. JNJ-678 was given at 6 hpi (Figure 1J) so that early events of RSV infection were not disrupted. JNJ-687 effectively prevented viral spread evident by the reduction in the number of RSV-infected cells and RSV RNA levels in both neonatal and adult ALI cultures compared to solvent control (Figure 1J). In addition, JNJ-678 treatment completely abrogated the difference in RSV viral RNA levels between neonatal and adult cultures at 1 dpi and 2 dpi (Figure 1J). Therefore, age affects the spread of RSV in human bronchial epithelium models.

### Age-related apoptotic cell death causes severe RSV infection in neonatal bronchial epithelium

RSV infection induces bronchial epithelial cell apoptosis in severe disease in infants^3,44^. Given that cell death is a means of viral spread^45^, we tested whether age affects apoptosis of RSV-infected ciliated cells and whether apoptosis contributes to viral spread. Employing cleaved caspase 3 (c-Casp-3) as a marker for apoptosis, we found negligible apoptotic cell death in neonatal and adult ALI cultures without infection (Figures 2A-2D). Following RSV infection, c-Casp-3 was barely detectable in adult ALI cultures by both antibody staining and Western blot (Figure 2A-2D), consistent with a lack of cytopathy in adult epithelial cells (Figure 1C)^29^. In contrast, RSV triggered significant apoptosis in neonatal ALI cultures (Figures 2A-2D). At 2 dpi, ∼25% of cells in neonatal ALI cultures were RSV F^+^ and nearly 80% of these were also c-Casp-3^+^ (Figures 2A and 2B). By 4 dpi, the percentage of RSV F^+^c-Casp-3^+^ cells in neonatal ALI cultures was reduced to ∼10% likely due to sloughing of RSV-infected cells (Figures 1C and S2C). We found an increased number of c-Casp-3^+^ cells below the apical surface between 2 dpi and 4 dpi that were not infected by RSV (Figure 2A). Apoptosis of bystander cells in RSV-infected neonatal ALI cultures may be induced by released inflammatory signals, such as TNF (Figure S3C). In addition to apoptosis, a small number of neonatal epithelial cells, including both RSV F^+^ and RSVF^-^ cells, also expressed necroptotic markers (Figure S5), which is consistent with a previous report that RSV infection can induce necroptosis^8^. Furthermore, despite similar viral RNA levels in JNJ-678-treated neonatal and adult ALI cultures at 2 dpi (Figure 1J), only neonatal cells were c-Casp-3^+^ (Figure S6A), which indicates that age-related apoptosis induced by RSV infection of neonatal ciliated cells is not caused by the disparity in viral replication and/or spread. Taken together, neonatal ciliated cells are predisposed to apoptosis following RSV infection, a response that is outgrown in adulthood.

**Figure 2.**
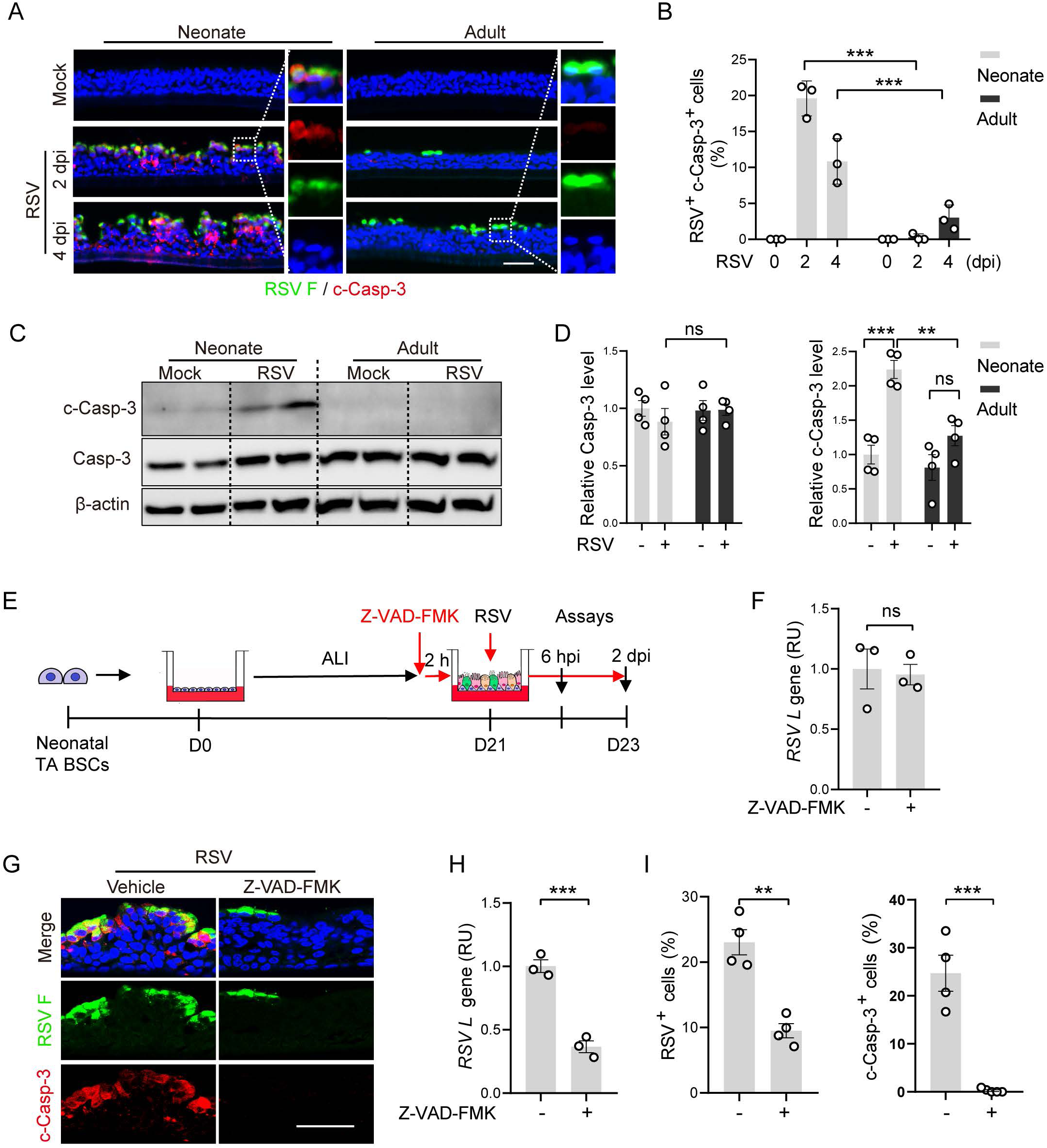
Age-related apoptotic cell death causes severe RSV infection in neonatal bronchial epithelium. (A) Representative double staining for RSV F and c-Casp-3 using sections of neonatal and adult ALI cultures (n=4 BSC lines per age group) at 2 and 4 dpi. (B) The relative abundance of RSV F^+^c-Casp-3^+^ cells quantified from double stained images. (C) Representative Western blot analyses of Caspase-3 (Casp-3) and c-Casp-3 at 2 dpi. β-actin was loading control. Each lane represents one BSC line. (D) Densitometry measurements of relative levels of Casp-3 and c-Casp-3 normalized to β-actin. (E) Schematic of Z-VAD-FMK treatment of neonatal ALI cultures (n=4 BSC lines). Z-VAD-FMK (40 μM) was applied in the bottom chamber 2 hours prior to RSV infection until 2 dpi. Assays were performed at 6 hpi and 2 dpi and shown in (F-I). (F) Relative levels of RSV *L* gene at 6 hpi by RT-qPCR. (G) Representative double staining for RSV F and c-Casp-3 at 2 dpi. (H) Relative levels of RSV *L* gene at 2 dpi by RT-qPCR. (I) The relative abundance of RSV F^+^ and c-Casp-3^+^ cells at 2 dpi quantified from double stained images. Each dot represents one BSC line. Bar graphs represent mean ± SEM. Statistical significance was calculated by two-way ANOVA followed by Sidak’s multiple comparison test in (B and D) and Student’s t-test (two-tailed) in (F, H, and I). **p<0.01, ***p<0.001, and ns, not significant. Scale bars, 50 μm.

To assess whether apoptosis facilitates RSV spread, we treated RSV infected neonatal ALI cultures with a caspase inhibitor, Z-VAD-FMK, starting 2 hours prior to infection (Figure 2E). At 6 hpi, the viral RNA level was comparable between solvent- and Z-VAD-FMK-treated neonatal ALI cultures (Figure 2F). Therefore, Z-VAD-FMK treatment has no effect on early events of RSV infection. At 2 dpi, Z-VAD-FMK treatment prevented neonatal epithelial cells from apoptosis and reduced the number of RSV-infected cells and viral RNA by more than 50% compared to solvent control (Figures 2G-2I). Z-VAD-FMK treatment also ameliorated mucous hyperplasia and decreased the level of inflammatory gene expression in RSV-infected neonatal ALI cultures (Figures S6B and S6C). Furthermore, treatment with a second caspase 3 inhibitor, Z-DEVD-FMK, achieved similar beneficial results (Figure S6D). These findings collectively show that apoptosis of RSV-infected ciliated cells promotes viral spread.

### Age affects STAT3 activation in ciliated cells following RSV infection

STAT3 is a transcriptional factor involved in innate immunity^46^ and plays a critical role in cell survival by promoting the transcription of anti-apoptotic target genes within the *BCL2* family^47–49^. To test whether STAT3 signaling in human bronchial epithelium was associated with age, we measured the levels of total STAT3 and activated STAT3 (p-STAT3^Y705^) in neonatal and adult ALI cultures. Western blot analyses showed similar levels of total STAT3 levels between the two age groups with and without RSV infection (Figure 3A). In contrast, the relative level of p-STAT3^Y705^ at baseline was ∼30% higher in adult ALI cultures compared to neonatal ALI cultures (Figures 3A and 3B). Following RSV infection at 2 dpi, only adult ALI cultures exhibited STAT3 activation with p-STAT3^Y705^ levels elevated above an already higher baseline to 70% more than neonatal cultures (Figures 3A and 3B).

**Figure 3.**
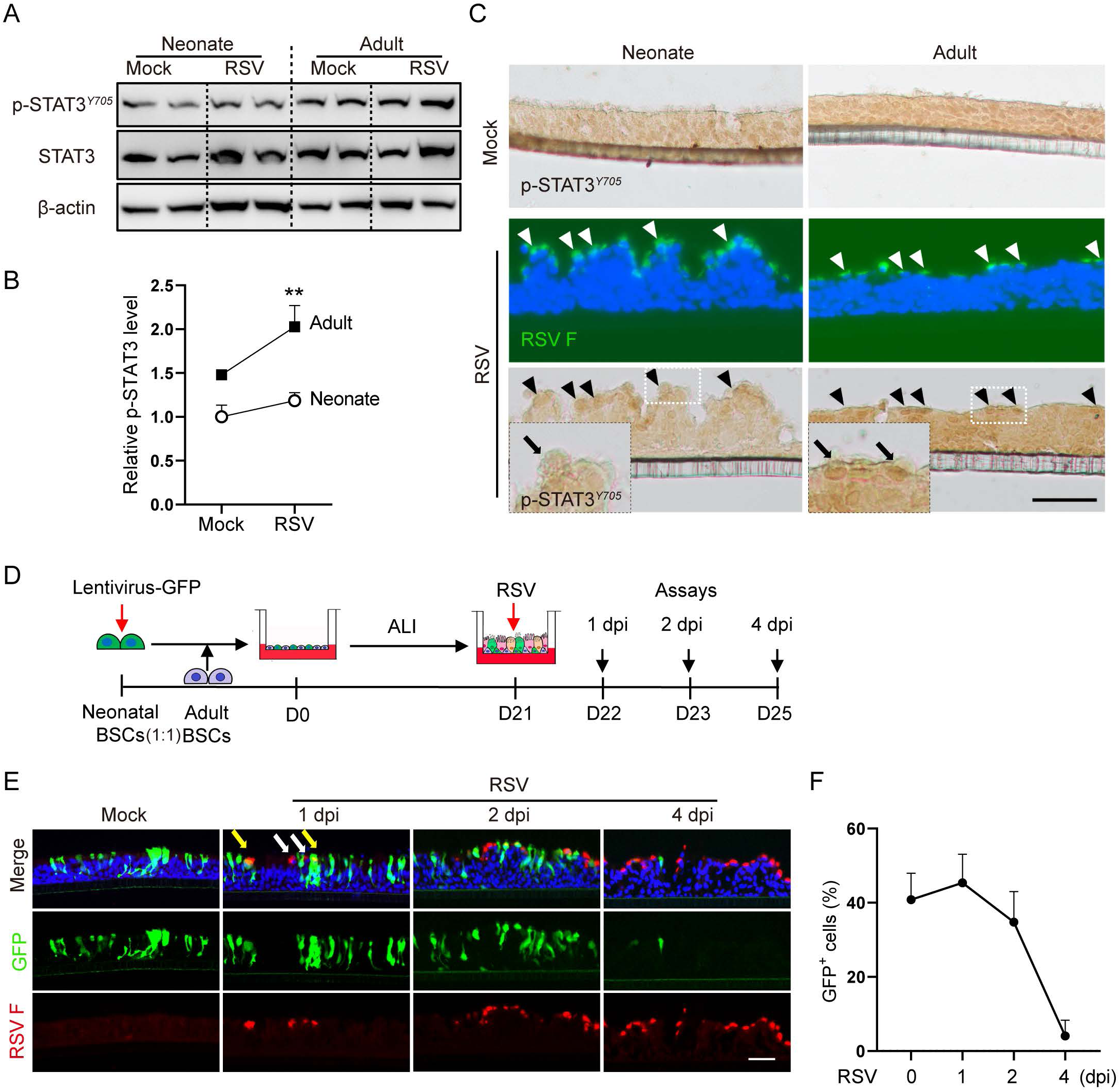
Age regulates STAT3 activation following RSV infection in ciliated cells. (A) Representative Western blot analyses of p-STAT3^Y705^ and STAT3 levels at 2 dpi in neonatal and adult ALI cultures (n=3 BSC lines per age group). β-Actin was loading control. Each lane represents one BSC line. (B) Densitometry measurements of the relative level of p-STAT3^Y705^ normalized to β-Actin. **p<0.01 by two-way ANOVA followed by Sidak’s multiple comparison test. (C) Representative double staining for RSV F (fluorescence) and p-STAT3^Y705^ (chromogenic). Arrowhead marks RSV F^+^ ciliated cells. Inserts show enlarged images of p-STAT3^Y705^ staining. Scale bar, 50 μm. (D) Schematic of RSV infection of a hybrid ALI culture. GFP-labelled neonatal BSCs (n=2 lines) were mixed with adult BSCs (n=2 lines) at a 1:1 ratio followed by differentiation in ALI. Hybrid ALI cultures were analyzed at 1, 2, and 4 dpi. (E) Representative staining for RSV F in hybrid ALI cultures. White arrows mark RSV F^+^GFP^-^ (adult) cells and yellow arrows mark RSV F^+^GFP^+^ (neonatal) cells. Scale bar, 50 μm. (F) The relative abundance of GFP^+^ cells in hybrid ALI cultures.

We performed antibody staining for p-STAT3^Y705^ to assess whether STAT3 activation in response to RSV infection was cell type-specific. Baseline nuclear p-STAT3^Y705^ was ubiquitously distributed in both neonatal and adult ALI cultures after extended chromogenic exposure (Figure 3C). At 2 dpi, nuclear p-STAT3 became readily detectable only in adult ciliated cells infected with RSV (Figure 3C), consistent with the increase in p-STAT3 levels in adult ALI cultures detected by Western blot (Figures 3A and 3B). Given age-related differences in the number of RSV-infected cells at 2 dpi (Figure 1E), we estimated that the p-STAT3^Y705^ level per RSV-infected ciliated cell in adult cultures was minimally 5-fold higher than that in neonatal cultures. Lastly, STAT3^Y705^ levels positively correlated with differences in BCL2 family gene expression between neonatal and adult ALI cultures at baseline and following RSV infection (Figure S7A).

To test whether STAT3 acts in a ciliated cell-autonomous manner to regulate apoptosis following RSV infection, we mixed neonatal and adult BSCs at a 1:1 ratio to generate a hybrid bronchial epithelium model in ALI (Figure 3D). Neonatal BSCs were pre-labelled by a GFP lentivirus to track their progenies (Figures 3D and 3E). GFP^+^ (neonatal) and GFP^-^ (adult) ciliated cells were found at an equal frequency in the apical surface of hybrid ALI cultures at day 21 and had similar rates of RSV infection at 1 dpi (Figures 3E and 3F). The number of RSV^+^GFP^+^ cells (neonatal) was decreased at 2 dpi and by 4 dpi, only a few GFP^+^ cells remained (Figures 3E and 3F). The kinetics of neonatal cell loss in the hybrid culture was similar to that in neonatal (only) ALI cultures following RSV infection (Figures 1C, 2A, and 2B). Therefore, the presence of adult epithelial cells imparted no protection from apoptosis to neonatal epithelial cells, which supports a cell-autonomous role of STAT3 in regulating apoptosis. Notably, at 4 dpi, compared to adult (only) ALI cultures in which ∼8% of epithelial cells were RSV^+^ (Figure 1E), hybrid cultures had up to 15% of RSV^+^GFP^-^ cells (adult) despite a significant loss of initially infected neonatal cells (Figures 3E and 3F). These findings provide further evidence in support of apoptosis-mediated viral spread from RSV-infected neonatal epithelial cells to neighboring adult cells.

### SARS-CoV-2 infection of human bronchial epithelium models is not age-related

After demonstrating that neonatal and adult bronchial epithelium models reproduce age-related RSV infection, we employed these models to test whether epithelial cell interaction with SARS-CoV-2 differs with age (Figure 4A). Antibody staining for SARS-CoV-2 nucleocapsid protein (N) showed that similar to RSV, SARS-CoV-2 also targeted ciliated cells in ALI cultures (Figures 4B and 4E)^23,24^. At 1 dpi, there was no difference in the number of SARS-CoV-2-N^+^ cells between neonatal and adult ALI cultures (Figures 4B-4C). The number of SARS-CoV-2-N^+^ cells increased significantly between 1 dpi and 3 dpi, which was similar between the two ages (Figures 4B-4D). In addition, we found no evidence for apoptotic cell death and STAT3 activation following SARS-CoV-2 infection in either age group (Figures 4E and 4F). Instead, nuclear p-STAT3^Y705^ was found in cells located just above the insert membrane that were likely the remaining basal cells (Figure 4E). Consistent with our findings *in vitro*, antibody staining of adjacent lung sections (4 µm apart) prepared from postmortem biopsy samples of fatal COVID-19 showed no c-Casp-3 or p-STAT3^Y705^ in SARS-CoV-2-N^+^ ciliated cells (Figure 4G). Notably, c-Casp-3 was found in non-epithelial cells (Figure 4G) and p-STAT3^Y705^ was shown to be localized to a subset of regenerating BSCs in these postmortem lung samples^35^. Taken together, SARS-CoV-2 infection of human bronchial epithelium is not related to age, ciliated cell apoptosis, or STAT3 regulation, which contrasts sharply with RSV infection.

**Figure 4.**
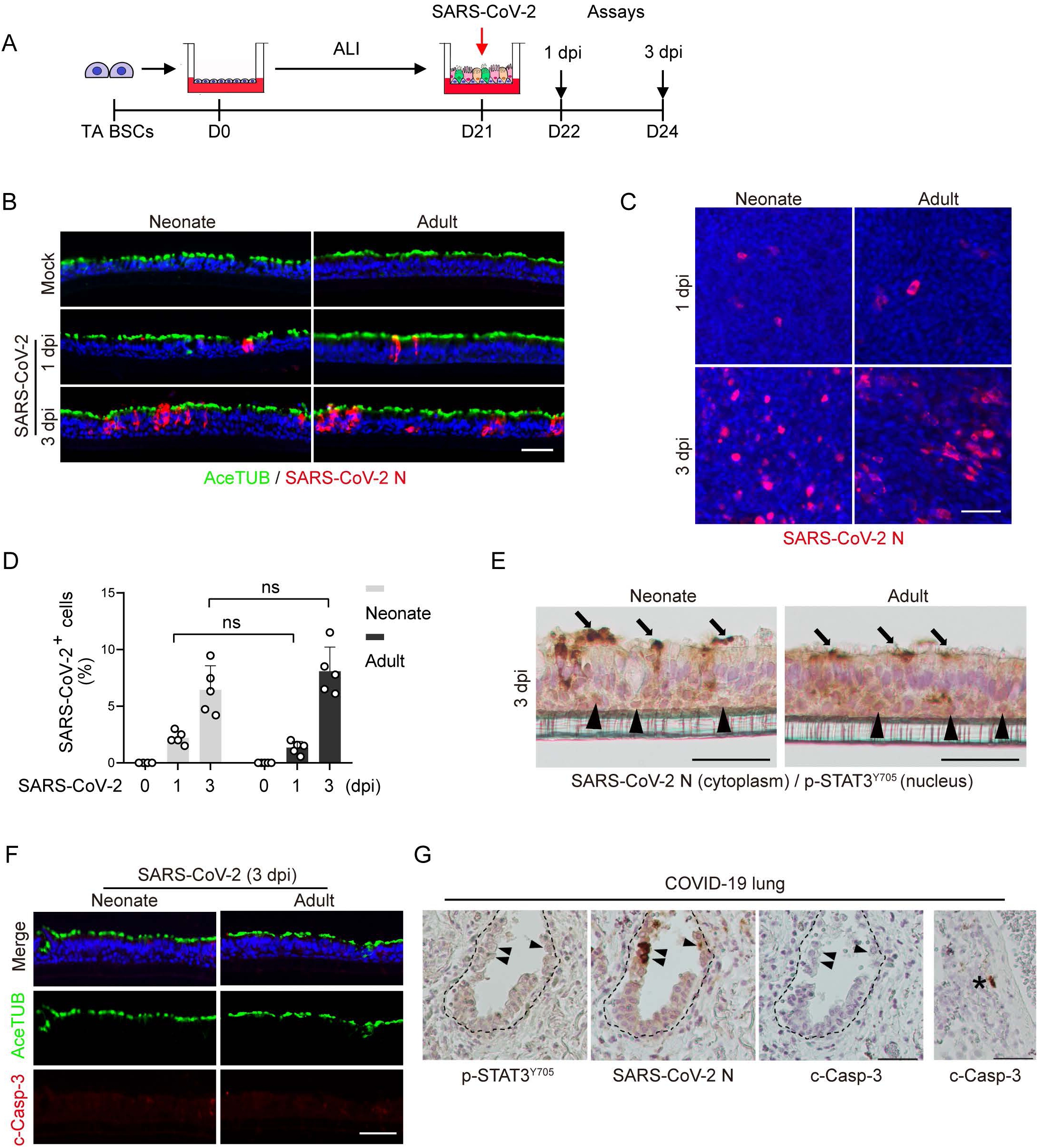
Age has no effect on SARS-CoV-2 infection of human bronchial epithelial cells. (A) Schematic of SARS-CoV-2 infection (MOI 3) of day 21 ALI cultures of neonatal and adult TA BSCs (n=2 BSC lines for each age group) followed by analyses at 1 and 3 dpi. (B) Representative double staining for AceTUB and SARS-CoV-2 N in cross sections of ALI cultures. (C) Representative whole-mount staining for SARS-CoV-2 N. (D) The relative abundance of SARS-CoV-2 N^+^ cells in neonatal and adult ALI cultures. (E) Representative chromogenic double staining for SARS-CoV-2 N (cytoplasm) and p-STAT3^Y705^ (nucleus) at 3 dpi. Arrows point to SARS-CoV-2 N^+^ ciliated cells. Arrowheads point to cells positive for nuclear p-STAT3^Y705^. (F) Representative double staining for c-Casp-3 and AceTUB at 3 dpi. (G) Representative staining for p-STAT3^Y705^, SARS-CoV-2 N, and c-Casp-3 using three adjacent lung sections (4µm apart) of post-mortem lung samples from fatal COVID-19 patients (n=5 donors). The dashed line marks basement membrane. Arrowheads mark SARS-CoV-2 N^+^ ciliated cells. Asterisk marks a c-Casp-3^+^ cell in lung parenchyma that was negative for SARS-CoV-2 N. Each data point (D) represents quantifications of one 20X image of one BSC line. Bar graph represents mean ± SEM. ns, not significant by two-way ANOVA followed by Sidak’s multiple comparison test. Similar results were observed in 2 neonatal and 2 adult BSC lines. Scale bars, 50 μm.

### RSV-infected ciliated cells can spread RSV after extrusion from neonatal bronchial epithelium model

Mucosal epithelium extrudes apoptotic cells to maintain the integrity of barrier function^50,51^. To test whether RSV exploited extrusion to spread the virus from apoptotic RSV-infected ciliated cells, we collected extruded cells in apical washes of neonatal ALI cultures at 2 dpi and 4 dpi (Figure 5A). The extruded cells were RSV-infected ciliated cells (Figure 5B) and the cell number increased by more than 3-fold between 2 dpi and 4 dpi in neonatal ALI cultures (Figure 5C). Almost all extruded cells were c-Casp3^+^, indicating ongoing apoptosis (Figure 5D). In contrast, few epithelial cells were extruded in RSV-infected adult ALI cultures (Figures 5B and 5C), consistent with a lack of apoptosis in these cultures (Figures 2A-2C)^29^. TUNEL staining of extruded neonatal cells for fragmented DNA, a marker of the final stage of apoptotic cell death, showed an increase in the abundance of terminally dying cells from less than 20% at 2 dpi to ∼60% at 4 dpi (Figure 5E). These findings are consistent with extrusion of epithelial cells during the early phase of apoptosis^52,53^. Our results also provide evidence that the completion of the process of apoptotic cell death in extruded RSV-infected neonatal ciliated cells likely takes more than 2 days, which provides a time window for these extruded cells to spread RSV.

**Figure 5.**
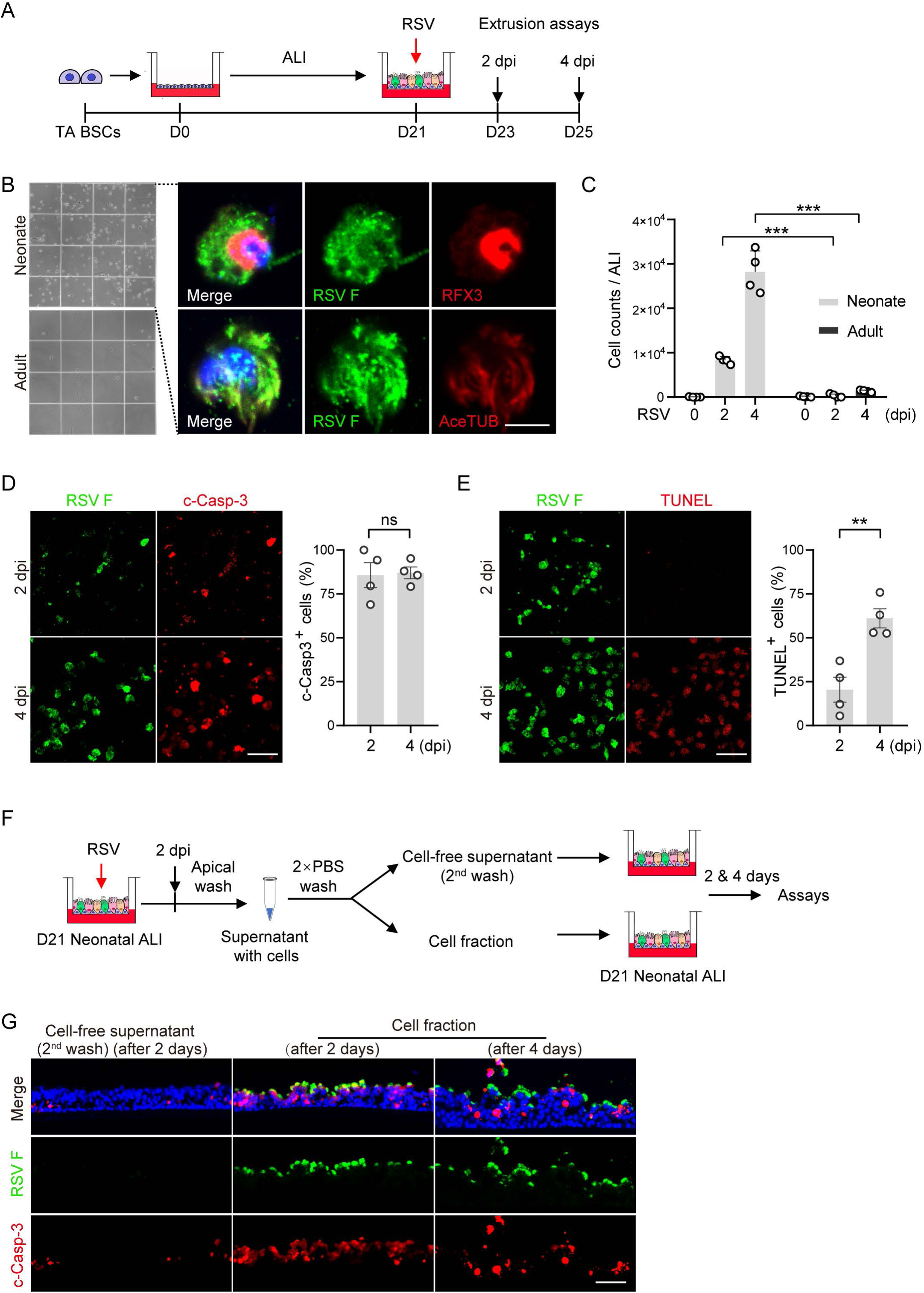
Cell extrusion in neonatal epithelium model following RSV infection can mediate viral spread. (A) Schematic of extrusion assays. Extruded cells were collected from apical washes of neonatal and adult ALI cultures (n=4 BSC lines per age group) at 2 and 4 dpi. Extruded cells were analyzed in (B-E). (B) Representative images of extruded cells on a cytometer (left panels) and double staining for RSV F and ciliated cell markers (RFX3 and AceTUB) (right panels). (C) Quantification of the number of extruded cells. (D) Representative double staining for RSV F and c-Casp-3 (left panels) and quantification of the relative abundance of c-Casp-3^+^ cells (right panel). (E) Representative double staining for RSV F and TUNEL (left panels) and quantification of the relative abundance of TUNEL^+^ cells (right panel). (F) Schematic of infection assay by extruded cells. Apical washes collected from RSV-infected neonatal ALI cultures at 2 dpi were centrifuged and cell pellets were washed 2 times. Cell fraction and cell-free supernatant from the 2^nd^ wash were applied apically to uninfected neonatal ALI cultures for 12 hours. The cultures were analyzed for RSV infection after 2 and 4 days in (G). (G) Representative double staining for RSV F and c-Casp-3 after treatment with cell fraction and cell-free supernatant. Each dot represents one BSC line. Bar graphs show mean ± SEM. ***p<0.001 calculated by two-way ANOVA followed by Sidak’s multiple comparison test in (C). **p<0.01 and ns, not significant by Student’s t-test (two-tailed) in (D and E). Scale bars, 5 μm in (B) and 50 μm in (D, E, and G).

To test whether extruded ciliated cells facilitate RSV spread, we collected the cell fraction from apical washes of neonatal ALI cultures at 2 dpi by centrifugation (Figure 5F). After two rounds of saline washes to remove extracellular virions, the cell pellet was resuspended in saline and cell-free supernatant from the second wash was collected (Figure 5F). The cell fraction applied apically to uninfected neonatal ALI cultures overnight was able to induce widespread RSV infection assayed after 2 and 4 days (Figures 5F and 5G). This was not due to carryover of already-secreted RSV virions because cell-free supernatant failed to establish infection (Figure 5G). Therefore, extruded, RSV-infected ciliated cells can spread RSV during the process of apoptotic cell death.

### STAT3 inhibition in adult bronchial epithelium model promotes apoptosis and worsens RSV infection

To test how age-related STAT3 activation is functionally connected to apoptosis and RSV viral spread, we inhibited STAT3 activity in adult ALI cultures using 3 different approaches. In the first approach, we treated adult ALI cultures with a specific STAT3 inhibitor, Stattic, 2 hours prior to RSV infection until 2 dpi (Figure 6A). The dose of Stattic (20 µM) was determined based on its efficacy in blocking the increase in p-STAT3^Y705^ levels in response to IL6 (Figure S8A). Of note, Stattic treatment had no effect on the expression of a RSV receptor known to be expressed by ciliated cells, IGF1R^54^ (Figure S8B) or barrier function of well-differentiated epithelial cells in adult ALI cultures^55^. At 2 dpi, Stattic treatment doubled the number of RSV-infected cells and the viral RNA level compared to solvent control (Figures 6B-6D). Stattic treatment during RSV infection also triggered apoptosis in ∼8% of adult epithelial cells in ALI compared to almost 0% in solvent control (Figures 6B and 6D). In a second approach, we employed a doxycycline (Dox)-inducible shRNA system to knockdown STAT3. Dox was administered from day 18 in ALI (Figures 6E and 6F), when multiple epithelial cell types are already generated^37,38^, to minimize the effect of STAT3 inhibition on ciliated cell differentiation^55,56^. STAT3 knockdown in adult ALI cultures resulted in almost 3 times more RSV-infected cells and induced apoptosis in ∼60% of RSV-infected cells (Figures 6G and 6H). In the third approach, we isolated bronchial BSCs from lung biopsy samples from healthy adult donors and an adult patient with Job syndrome harboring a hypomorphic STAT3-S560del mutation^55^ (Figure S9A). As previously reported^55^, STAT3-S560del BSCs had a reduced capacity to differentiate into ciliated cells compared to healthy biopsy BSCs (Figures S9B and S9C). However, despite fewer target cells for RSV infection, STAT3-S560del adult ALI cultures had more RSV F^+^c-Caps-3^+^ cells than healthy adult controls (Figures S9D and S9E). Lastly, the approaches to inhibit STAT3 in adult ALI cultures reduced *BCL2* gene expression (Figures S7B and S7C), consistent with *BCL2* being a direct target gene of activated STAT3^47,48^. Taken together, STAT3 activation in adult epithelial cells following RSV infection is required to prevent apoptosis and reduce severity of infection.

**Figure 6.**
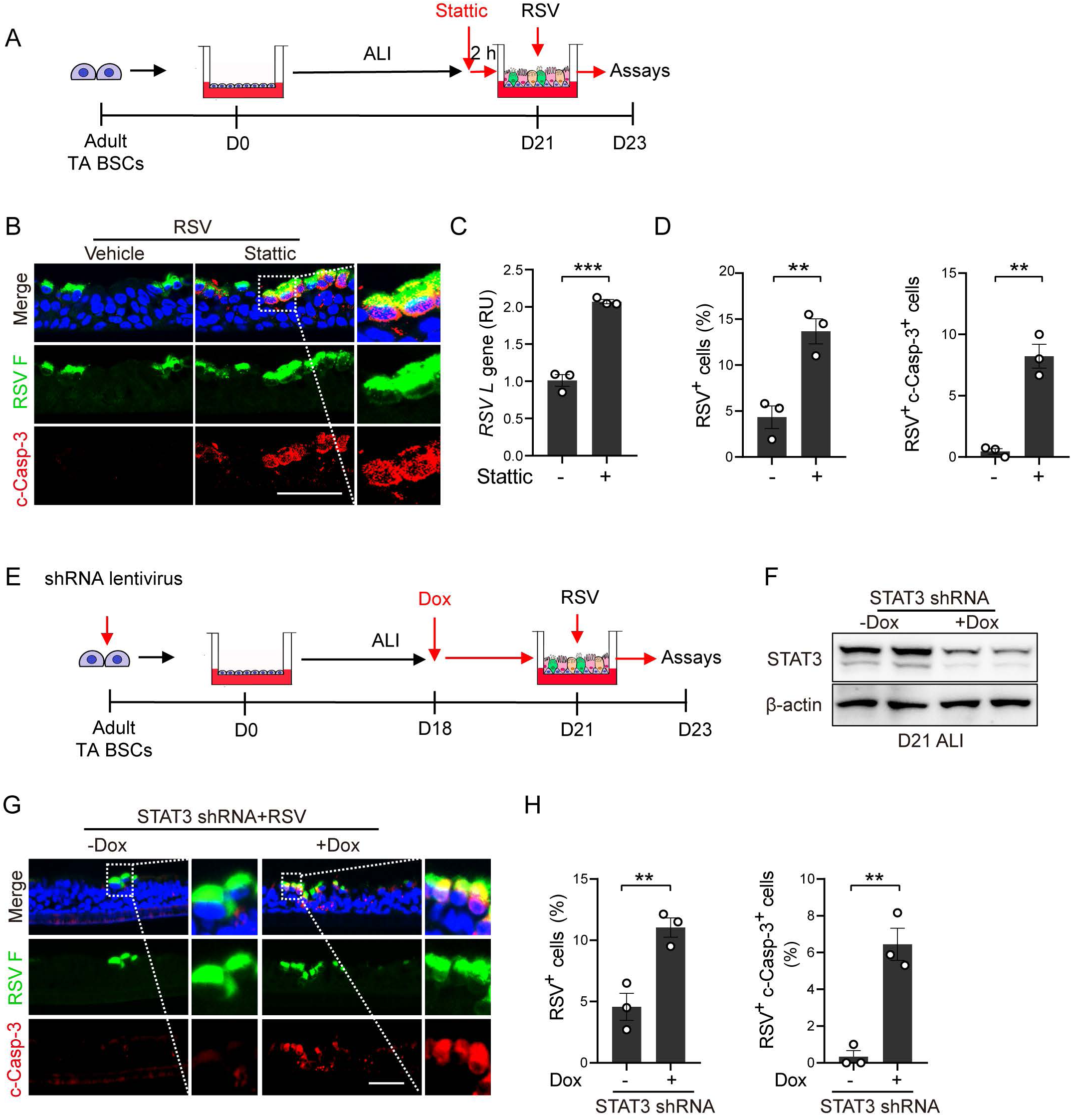
Blockade of active STAT3 in RSV-infected adult bronchial epithelium model worsens infection and promotes apoptosis. (A) Schematic of Stattic (20 μM) treatment. Stattic was applied in the bottom chamber of adult ALI cultures (n=3 BSC lines) 2 hours prior to RSV infection until 2 dpi. Results were shown in (B-D). (B) Representative double staining for RSV F and c-Casp-3. (C) The relative level of RSV *L* gene by RT-qPCR. (D) The relative abundance of RSV F^+^ and RSV F^+^c-Casp-3^+^ cells. (E) Schematic of STAT3 knockdown assay using an inducible lenti-shRNA system. ALI cultures of lentivirus transduced adult BSCs (n=3 lines) were treated with Dox (500 ng/mL) in the bottom chamber from day 18 and infected with RSV at day 21. Assays were performed 2 dpi and results were shown in (F-H). (F) Representative Western blot analyses to assess STAT3 knockdown efficiency. β-actin was loading control. Each lane represents one BSC line. (G) Representative double staining for RSV F and c-Casp-3. (H) Quantification of the relative abundance of RSV F^+^ cells and RSV F^+^c-Casp-3^+^ cells. Each dot represents one BSC line. **p<0.01 and ***p<0.001 calculated by Student’s t-test (two-tailed). Scale bars, 50 μm.

### STAT3 activation in neonatal bronchial epithelial cells reduces severity of RSV infection

To test whether augmenting STAT3 activation in neonatal epithelial cells could mitigate severe RSV infection, neonatal ALI cultures were treated with IL6, a cardinal activator of STAT3, at day 18 in ALI until 2 dpi (Figure 7A). The timing and the duration of IL6 treatment were determined based on the efficacy of increasing p-STAT3 levels and *BCL2* gene expression (Figures 7B, 7C, and S7D) and preserving mucociliary differentiation in neonatal ALI cultures (Figure S10). We showed that IL6 treatment had no effect on early events of RSV infection assayed at 6 hpi (Figure 7D). By 2 dpi, IL6-treated neonatal ALI cultures had a significant reduction in the number of RSV-infected and apoptotic ciliated cells, essentially resembling RSV-infected adult ALI cultures (Figures 7E and 7F, compared to Figures 1E and 2B). The beneficial effect of IL6 was blunted by overlapping Stattic blockade during RSV infection (Figures 7A, 7E, and 7F), indicating STAT3 as the critical mediator of IL6 treatment. Taken together, employing IL6 treatment as a proof-of-principle approach to boost STAT3 activity in neonatal bronchial epithelium model, our findings indicate that insufficient STAT3 activation is a major cause of severe RSV infection in neonatal bronchial epithelium.

**Figure 7.**
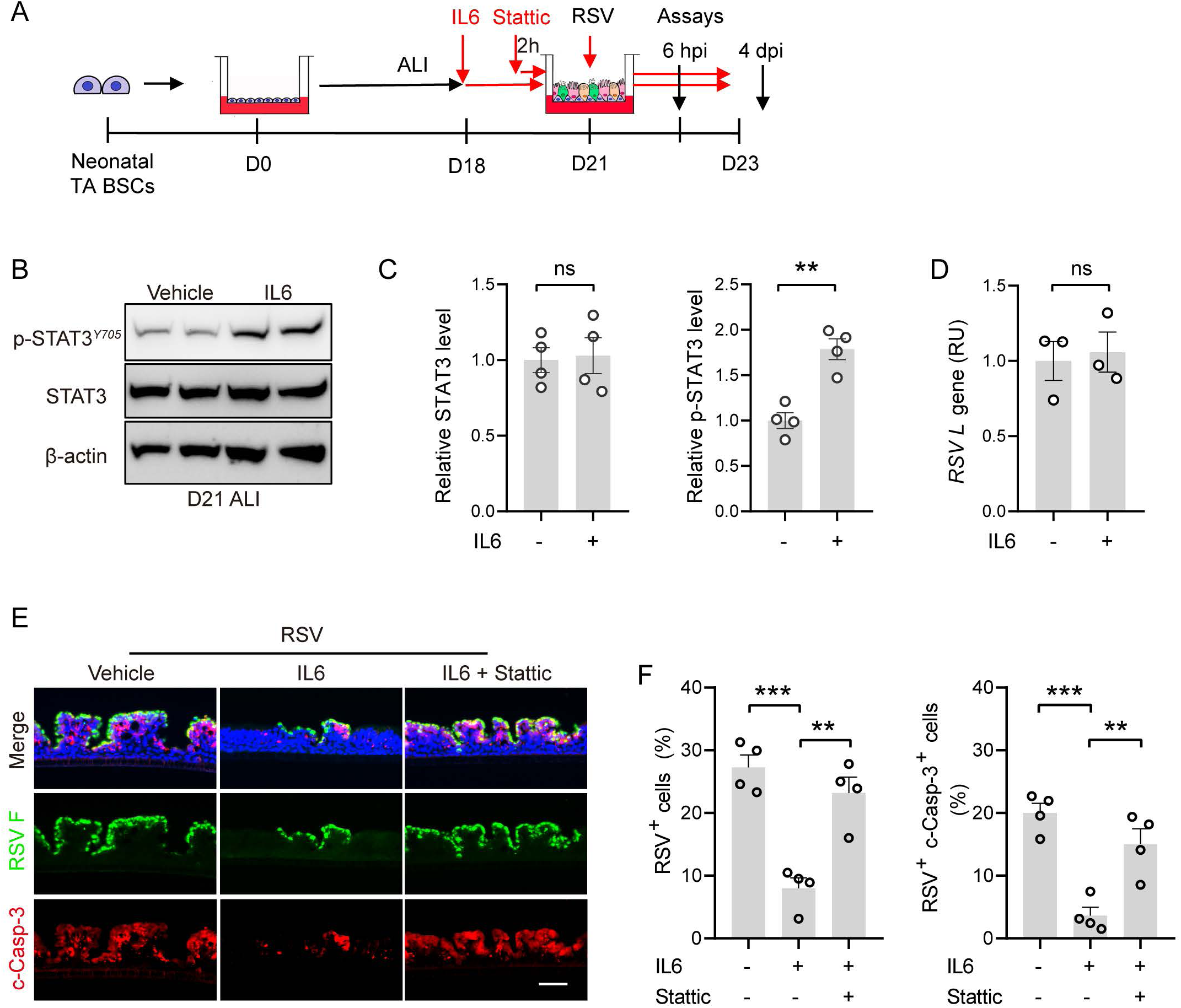
STAT3 activation in neonatal bronchial epithelial cells reduces severity of RSV infection. (A) Schematic of IL6 treatment assay. Neonatal cultures were treated with IL6 (50 ng/mL) in the bottom chamber at D18 in ALI with and without Stattic (20 μM) applied 2 hours prior to RSV infection. ALI cultures were assayed at 6 hpi and 2 dpi. (B) Representative Western blot analyses for levels of p-STAT3^Y705^ and STAT3. β-actin was loading control. Each lane represents one neonatal BSC line. (C) Densitometry measurement of relative levels of p-STAT3^Y705^ and STAT3. (D) The relative level of RSV *L gene* at 6 hpi by RT-q-PCR. (E) Representative double staining for RSV F and c-Casp-3. (F) The relative abundance of RSV F^+^ cells and RSV^+^c-Casp-3^+^ cells. Each dot represents one BSC line. Bar graphs show mean ± SEM. **p<0.01, ***p<0.001, and ns, not significant calculated by Student’s t-test (two-tailed) in (C and D) and **p<0.01 by one-way ANOVA followed Tukey’s multiple comparison test in (F). Scale bar, 50 μm.

## DISCUSSION

Leveraging an established method of TA BSC derivation, we generated well-differentiated human bronchial epithelial cells of neonates and adults to investigate how age affects epithelial cell infection by RSV and contributes to susceptibility to severe RSV disease in infants. Our study shows that RSV exploits age-related, insufficient STAT3 signaling in neonates to trigger apoptosis of infected ciliated cells, and that extruded apoptotic cells serve as a vehicle to facilitate viral spread. These findings provide evidence in support of the hypothesis that shredding of RSV-infected epithelial cells contributes to spread of infection as well as airway obstruction in severe RSV disease in infants.

RSV is equipped with additional virus-specific factors to capitalize on susceptibility to apoptotic cell death at an early age to cause severe infection. For example, RSV infection induces ciliary dyskinesia and mucus hyperplasia in infants to impair mucociliary clearance^4,40,41^. RSV non-structural proteins, including NS1 and NS2, also suppress premature apoptosis to promote viral replication^57^. Furthermore, RSV sensitizes infected cells to apoptosis activated by tumor necrosis factor-related apoptosis-inducing ligand^58^. These RSV-derived factors can potentially increase the number of apoptotic cells and prolong the time window of virion production and secretion from already-infected and extruded ciliated cells to spread RSV in bronchial epithelium.

Our study identifies STAT3 as a critical mediator of age-related severity of RSV infection of human bronchial epithelial cells. STAT3 is a multi-functional transcription factor that regulates the expression of genes involved in several biological processes such as inflammation, proliferation, and survival^59^. A variety of cytokines and growth factors, including IL6, IL22, and EGF^60^, can activate STAT3. In our study, we show that well-differentiated, adult bronchial epithelial cells secret higher levels of EGF (Figure S3E), which may explain higher levels of STAT3 activation in adults at baseline. Signals that further activate STAT3 in ciliated cells following RSV infection have yet to be identified. In addition, whether *BCL2* family genes are downstream mediators of STAT3 to regulate apoptotic cell death warrants future investigation. We show that complete inhibition of apoptosis in neonatal ALI model fails to completely reduce RSV viral RNA to similar, low levels found in the adult ALI model. Therefore, in addition to STAT3 regulation of apoptotic cell death, other age-related mechanisms are likely involved in determining severity of epithelial infection by RSV.

Mechanisms that regulate age-related STAT3 activation in ciliated cells following RSV infection remain to be identified. Because all differentiated epithelial cells are generated from BSCs in our model, BSCs must retain developmental and age-specific memory that is transmitted to their progeny, including ciliated cells, to mediate age-related responses to RSV. Our transcriptomic analyses validate the impact of age on BSCs. In support of our findings, a separate study reports no age-related difference in cellular composition in pediatric and adult bronchial epithelium *in vivo*; however, laser capture-microdissected whole epithelium shows changes in gene expression with age^39^. In addition to further characterization of BSCs at different ages, epigenetic and gene expression assays to identify age-related differences in their progeny differentiated ciliated cells are also warranted. The TA BSC-based epithelium model can be applied to patients across the age spectrum to investigate the transition from susceptibility to relative protection and back to susceptibility in the elderly. This approach would allow even deeper understanding how disease susceptibility is regulated at the RSV-host epithelium interface and whether additional genetic and environmental factors that shift this age of transition.

In contrast to RSV infection, SARS-CoV-2 infection has no effect on apoptosis or STAT3 activation. We also found that SARS-CoV-2 similarly infected neonatal and adult bronchial epithelial cell models. Consistent with our observation, there is no difference in SARS-CoV-2 receptor expression in human lungs between children and adults^61^. Based on these findings, we conclude that infection of human bronchial epithelial cells with SARS-CoV-2 is not affected by the age of the host. The observed differences in COVID-19 disease severity between adults and children are likely caused by an unchecked inflammatory response in adult patients^62–65^ that is different from the immune response in SARS-CoV-2-infected infants and young children^66,67^. The comparison between RSV and SARS-CoV-2 infection of human epithelium models highlights virus-specific mechanisms underlying age-related predilection for severe disease.

In summary, age affects bronchial epithelial cell interaction with RSV and contributes to severity of RSV infection in infants. Our study identifies prophylactic activities of STAT3 against severe RSV infection in neonatal bronchial epithelium by preventing apoptotic cell death and subsequent viral spread. Considering that respiratory viral infection in different areas of the lung is heterogenous, STAT3 activation may produce therapeutic benefits by reducing the spread of RSV to uninfected lung areas in infants with active RSV disease. Our neonatal bronchial epithelium model can be employed to identify additional molecular mechanisms of age-related RSV infection as potential therapeutic targets to treat severe RSV disease in infants.

## ACKNOWLEDGEMENTS

This work was supported by NIH grants to HM (R21AR080778), PHL (R21AI156597), JQ (R01HL152293, R01HL159675), XA (R21AI173494), and Fast Grants and Evergrande MassCPR to EM and funds from the Department of Pediatrics at MGH for Lung Cell Bank to XA.

## AUTHOR CONTRIBUTIONS

C.Z. performed RSV infection experiments. W.W. analyzed the transcriptomes of neonatal and adult BSCs. Y. B. performed human cytokine array. G.A. derived TA BSCs. H.M. provided BSCs from a patient with Job syndrome. J.O., A.J.H., and E.M. performed SARS-CoV-2 infection experiment. Y.F. and J.Q. contributed postmortem lung samples from patients with COVID-19. R.F. guided RSV infection experiments. X.A. performed antibody staining of postmortem lung samples. X.A. and P.H.L. conceived the study. C.Z. and X.A. wrote the manuscript. P.H.L. edited the manuscript. All the authors read and commented on the manuscript.

## STAR METHODS

### KEY RESOURCES TABLE

| REAGENT or RESOURCE | SOURCE | IDENTIFIER |
| --- | --- | --- |
| **Antibodies** |  |  |
| Mouse anti-RSV Fusion protein | Bio-Rad | Cat# MCA490, RRID: AB_2231368 |
| Rabbit anti-TP63 | Cell Signaling Technology | Cat# 13109, RRID: AB_2637091 |
| Rabbit anti-SCGB1A1 | Sigma-Aldrich | Cat# HPA031828, RRID: AB_2674061 |
| Mouse anti-MUC5AC | Thermo Fisher Scientific | Cat# MA5-12178, RRID: AB_10978001 |
| Rabbit anti-RFX3 | Sigma-Aldrich | Cat# HPA035689, RRID: AB_10671224 |
| Mouse anti-AceTUB | Sigma-Aldrich | Cat# T6793, RRID: AB_477585 |
| Rabbit anti-Cleaved Caspase-3 (Asp175) | Cell Signaling Technology | Cat# 9661, RRID: AB_2341188 |
| Rabbit anti-STAT3 | Cell Signaling Technology | Cat# 12640, RRID: AB_2629499 |
| Rabbit anti-phospho-Stat3 (Tyr705) | Cell Signaling Technology | Cat# 9145, RRID: AB_2491009 |
| Rabbit anti-phospho-RIP (Ser166) (D8I3A) | Cell Signaling Technology | Cat# 44590S, RRID: AB_2799268 |
| Rabbit anti-SARS-CoV nucleocapsid protein (N) | Rockland Immunochemicals | Cat# 200-401-A50, RRID: AB_828403 |
| Rabbit anti-ZO1 | Abcam | Cat# ab216880, RRID: AB_2909434 |
| Mouse anti-β-actin | Sigma-Aldrich | Cat# A5441; RRID: AB_476744 |
| Goat anti-rabbit IgG (H+L) AF594 | Thermo Fisher Scientific | Cat# A-11037, RRID: AB_2534095 |
| Goat anti-mouse IgG (H+L) AF488 | Thermo Fisher Scientific | Cat# A-11029, RRID: AB_2534088 |
| Goat polyclonal anti-rabbit, HRP | Santa Cruz Biotechnology | Cat# sc-2004; RRID: AB_631746 |
| Goat polyclonal anti-mouse, HRP | BD Biosciences | Cat# 554002; RRID: AB_395198 |
| **Virus strains** |  |  |
| RSV A2 | ATCC | VR-1540 |
| RSV B WV/14617/85 | ATCC | VR-1400 |
| SARS-CoV-2 USA_WA1/2020 | Kindly provided by CDC's Principal Investigator Natalie Thornburg and the World Reference Center for Emerging Viruses and Arboviruses (WRCEVA) | N/A |
| **Biological Samples** |  |  |
| Hep-2 cell line | ATCC | CCL-23 |
| Vero E6 cell line | ATCC | CRL-1586 |
| TA BSCs, see Table S1 |  |  |
| **Chemicals and recombinant proteins** |  |  |
| Stattic | Tocris | Cat# 2798 |
| Z-VAD-FMK | Selleck Chemicals | Cat# S7023 |
| Z-DEVD-FMK | ApexBio | Cat# A1920 |
| JNJ-678 | MedChem Express | Cat# HY-112180 |
| IL6 (human) Recombinant Protein | Thermo Scientific | Cat# RP8619 |
| Hoechst dye | Thermo Fisher Scientific | Cat# H3570 |
| Y-27632 2HCl | Selleck Chemicals | Cat# S1049 |
| Rapamycin 99% | Fisher Scientific | Cat# R-5000 |
| CHIR 99021 | Tocris Bioscience | Cat# 4423 |
| A83-01 | Tocris Bioscience | Cat# 2939 |
| Doxycycline hyclate, $\geq 98\%$ (HPLC) | Sigma-Aldrich | Cat# D9891-25G |
| Lipofectamine 3000 Transfection Reagent | Thermo Scientific | Cat# L3000015 |
| Normocin | InvivoGen | Cat# ant-nr-1 |
| Cell Lysis Buffer | Cell Signaling Technology | Cat# 9803S |
| Fluoromount-G | Southern Biotech | Cat# 0100-01 |
| Phosphatase inhibitor cocktails | EMD Millipore | Cat# 524627-1EA |
| Protease inhibitor | Sigma-Aldrich | Cat# 11697698001 |
| **Critical Commercial Assays** |  |  |
| Standard ABC kit | Vector Laboratories | Cat#PK-6100 |
| Hematoxylin QS | Vector Laboratories | Cat#H-3404 |
| DAB Peroxidase Substrate Kit | Vector Laboratories | Cat# SK-4100 |
| RNeasy kit | QIAGEN | Cat# 74106 |
| CF594-TUNEL Apoptosis Assay Kit | Biotium | Cat# 30064 |
| Superscript III Reverse Transcriptase Kit | Thermo Fisher | Cat# 18080-044 |
| SYBR Green PCR Master Mix | Thermo Fisher Scientific | Cat# 4367659 |
| SuperSignal West Dura Extended Duration Substrate | Thermo Fisher Scientific | Cat# 34075 |
| Human XL Cytokine Array Kit | R&D Systems | Cat# ARY022B |
| Small Airway Epithelial Cell Growth Medium (SAGM) | PromoCell | Cat# C-21070 |
| PneumaCult-ALI Medium | STEMCELL Technologies | Cat# 05001 |
| **Oligonucleotides** |  |  |
| Primers for qPCR and shRNA, see Table S2 | IDT | N/A |
| **Deposited data** |  |  |
| Bulk RNA-seq datasets of adult TA BSCs | Bankoti et al. <sup>35</sup> and this paper | GSE212412, GSE242397 |
| Bulk RNA-seq dataset of neonatal TA BSCs | Wagner et al. <sup>34</sup> and this paper | GSE211790, GSE242397 |
| **Software and Algorithms** |  |  |
| GraphPad Prism 8 | GraphPad Software | https://www.graphpad.com |
| ImageJ 1.49v | ImageJ | https://imagej.nih.gov |
| NIS-Elements (D4.30.01) | NIS-Elements | https://www.microscope.healthcare.nikon.com |
| R (1.30.0) | R Project | https://www.r-project.org |

## RESOURCE AVAILABILITY

### Lead contact

Further information and requests for resources and reagents should be directed to and will be fulfilled by the lead contact, Xingbin Ai.

### Materials Availability

Neonatal and neonatal TA BSCs are available upon request.

### Data and Code Availability

This paper analyzes new and preexisting bulk RNA-seq datasets of adult BSCs and neonatal BSCs. The accession numbers for the datasets are listed in the Key resources table.

## EXPERIMENTAL MODEL AND SUBJECT DETAILS

### Histological human lung tissue sections

Paraffin sections of healthy lung samples were prepared from de-identified donor lungs (0-69 years of age) purchased from International Institute for the Advancement of Medicine (IIAM). Four adult lungs (1 female and 3 male donors of 43-69 years of age) and 3 lungs from children (1 female and 2 males of 0-12 months of age) were used in this study. The donors had no previously known lung diseases. Because the experiment involves no intervention or interaction with living individuals, this project is deemed non-human subject research by the Institutional Review Board at MGH. Postmortem lung sections from deceased adult patients with COVID-19 were provided by Dr. Jianwen Que at Columbia University and described in our previous study^35^.

### BSC isolation and expansion

BSC derivation from TA samples was described previously^32^. Briefly, TA samples were collected via in-line suctioning from intubated neonatal and adult patients at the Intensive Care Unit (ICU), MGH under an approved IRB protocol (No. 2019P003296). Patients were intubated due to cardiogenic and neurogenic respiratory failure. BSCs in TA samples were expanded in Small Airway Epithelial Cell Growth Medium (SAGM, PromoCell, Cat# C-21070 in the presence of SMAD/ROCK/mTOR inhibitors^32–34^. BSCs of one patient with Job syndrome were isolated from fresh discarded surgical specimens under an approved IRB protocol (No. 2017P001479)^55^ and expanded similarly as TA BSCs.

### RSV and SARS-CoV-2 preparation

RSV strain A2 (VR-1540) and RSV strain B (WV/14617/85) were purchased from ATCC (VR-1400). RSV was propagated in Hep-2 cell line (ATCC CCL-23) as previously described (5). Briefly, Hep-2 cells were cultured in Opti-MEM (Gibco) supplemented with 2% fetal bovine serum (FBS) (Gibco) and 1% GlutaMAX Supplement (Gibco). At 80% confluency, Hep-2 cells were infected with RSV in complete medium at a multiplicity of infection (MOI) of 0.1 for 1 h at 37°C. When high cytopathic effects were observed (∼5 days post infection) in culture, cells were scraped into the medium, cell debris was separated by centrifugation, and the supernatant was aliquoted and stored at -80°C. RSV virus titers were measured via plaque assay as previously described^68^.

SARS-CoV-2 stocks [isolate USA_WA1/2020, provided Dr. Natalie Thornburg at CDC and the World Reference Center for Emerging Viruses and Arboviruses (WRCEVA)] were propagated in Vero E6 cells (ATCC CRL-1586) cultured in Dulbecco’s modified Eagle’s medium (DMEM) supplemented with 2% FBS, penicillin (50 U/mL), and streptomycin (50 mg/mL). To remove confounding cytokines and other factors, viral stocks were purified by ultracentrifugation through a 20% sucrose cushion at 80,000 g for 2 h at 4°C as previously described^69^. SARS-CoV-2 titers were determined in Vero E6 cells by tissue culture infectious dose 50 (TCID50) assay. All work with SARS-CoV-2 was performed in the biosafety level 4 (BSL-4) facility of the National Emerging Infectious Diseases Laboratories at Boston University following approved standard operating procedures.

## METHOD DETAILS

### Transcriptome Analysis of BSCs by bulk RNA-seq

Bulk RNA-seq service was provided by Active Motif (Carlsbad, CA, USA). As previously described ^34^, RNA (2 μg per sample) extracted from 1×10^6^ BSCs was performed using the Qiagen RNA RNeasy Kit (Cat#74104, Qiagen) followed by library preparation with the Illumina TruSeq V2 Kit (Cat#20020594, Illumina). Libraries were sequenced on Illumina NextSeq 500 as paired-end 42-nt reads and the reads were subsequently mapped to the human hg38 reference genome using STAR algorithm version 2.6.0a. Only genes with an average count above 2 were analyzed. The DESeq2 R package (1.30.0) was used to perform differential expression and principal component analyses. Genes with an adjusted p value less than 0.05 (padj<0.05) were considered as differentially expressed genes (DEGs).

### Mucociliary differentiation of BSCs in ALI

As previously described^32^, 2×10^5^ BSCs were seeded onto the 6.5 mm Transwell with 0.4 µm Pore Polyester Membrane Insert (Cat# 3470, Corning) that was precoated with 804G-conditioned medium. Differentiation in PneumaCult medium (Cat# 05001, StemCell Technology) started after removal of medium from the top chamber and completed after 21 days in ALI. Medium was changed every 2 days. At day 21 in ALI, cultures were processed for antibody staining and viral infection assay. For hybrid ALI culture experiments, neonatal BSCs were infected with a GFP lentivirus followed by treatment with puromycin (1 μg/mL) for 3 days to select for GFP-transduced BSCs. GFP^+^ neonatal BSCs were then mixed with adult BSCs at a 1:1 ratio and subjected to differentiation in ALI. For Dox-inducible STAT3 knockdown experiments in ALI cultures, after screening with puromycin (1 μg/mL) for 3 days, BSCs were subjected to differentiation in ALI culture. At 18 days of ALI differentiation, the cultures were treated with vehicle or Dox (500 ng/ml) in the basal medium. The knockdown efficiency and infection were performed at 21 days of differentiation. Lentivirus was generated from 293T cells using Lipofectamine 3000 Transfection Reagent (Cat#L3000015, Thermo Scientific).

### Infection of ALI cultures with RSV and SARS-CoV-2

Frozen aliquots of the virus were thawed and diluted in PBS to a designated titer immediately before use. Prior to infection, the apical surface of ALI cultures was washed 2 times with 200 μL PBS to remove the mucus. Viral suspension (100-200 µL) was then applied to the apical surface for 1 h at 37°C, 5% CO2, followed by 3 washes with 200 μL PBS to remove unbound viral particles. ALI cultures were analyzed at multiple time points after infection. For RSV infection experiments, the cytopathic effect of virus-infected ALI cultures was monitored daily by bright field microscopy. Apical washes (200 μL PBS for 20 min at 37 °C) were collected to evaluate apoptosis of detached cells and viral spreading from apoptotic cells, and the medium in the basal chamber was collected and stored at −80 °C until cytokine profiling using a kit following manufacture’s’ protocol (Cat# ARY022B, R&D Systems). At the end of infection experiments, RSV infected-epithelial cells on the insert were either fixed for 15 min in 4% paraformaldehyde/PBS for immunocytochemistry or collected in lysis buffer for protein or RNA assays. Following approved inactivation procedures, SARS-CoV-2 infected cells were fixed for at least 6 hours with 4% paraformaldehyde and removed from the maximum containment laboratory for further analysis.

### Antibody staining of ALI cultures and quantification

Mock or virus-infected ALI cultures were fixed for 15 min (RSV) or at least 6 hours (SARS-CoV-2) in 4% paraformaldehyde/PBS before they were processed for cryosectioning at a thickness of 10 μm or whole-mount staining. Sections or whole-mount ALI cultures were immune-stained according to a standard protocol using primary antibodies and secondary antibodies listed in the antibody table. To identify different epithelial cell types in ALI cultures, sections were stained for ciliated cells (rabbit anti-RFX3, 1:100, Cat# HPA035689,Sigma-Aldrich, and mouse anti-AceTUB, 1:100, Cat# T6793, Sigma-Aldrich), club cells (rabbit anti-SCGB1A1, 1:100, Cat# HPA031828, Sigma-Aldrich), goblet cells (mouse anti-MUC5AC, 1:100, Cat# MA5-12178, Thermo Fisher Scientific), and BSCs (rabbit anti-TP63, 1:100, Cat# 13109, Cell Signaling Technology). Additional antibodies used for the infected ALI cultures include: mouse anti-RSV Fusion protein (1:100, Cat# MCA490, Bio-Rad), rabbit anti-Cleaved Caspase-3 (Asp175) (1:100, Cat# 9661, Cell Signaling Technology), rabbit anti-phospho-RIP (Ser166) (D8I3A) (1:100, Cat# 44590S, Cell Signaling Technology), rabbit anti-SARS-CoV nucleocapsid protein (N) antibody that cross-reacts with SARS-CoV-2 N (1:2000, Cat# 200-401-A50, Rockland Immunochemicals), rabbit anti-ZO1(1:100, Cat# ab216880, Abcam), and rabbit Phospho-Stat3 (Tyr705) (1:25, Cat# 9145, Cell Signaling Technology). Nuclei were stained with Hoechst dye (1:500, ThermoFisher, Cat# H3570). The following secondary antibodies were used accordingly: goat anti-rabbit Alexa Fluor 594 (IgG, 1:200, Cat#A-11037, Thermo Scientific) and goat anti-mouse Alexa Fluor 488 (IgG, 1:200, Cat#A-11029, Thermo Scientific).

For quantification, 4-6 randomly selected and non-overlapping fields in cross sections collected from ALI cultures of one BSC lines in 2-3 independent experiments were imaged at 20X. The total number of epithelial cells per image was counted based on the nuclei countered stained with Hoechst dye. More than 1000 nuclei were quantified for each ALI culture. Epithelial cells labelled with specific markers in the same image were counted and reported as the percentage of the epithelial cell population.

### Quantitative Real Time PCR (qPCR)

Total RNA was extracted from ALI cultures using a RNeasy kit (Qiagen, 74106) followed by reverse transcription using Superscript III Reverse Transcriptase (Thermo Fisher Scientific, 18080-044). Real time PCR was performed with SYBR Green Mix (Cat#4367659, Thermo Scientific) using CFX96 real-time system (Bio-Rad). Relative gene expression was measured by normalizing to *GAPDH* using ΔΔCt (cycle threshold difference). All primers were listed in Table S2.

### Western Blot

ALI protein was harvested in 1X cell lysis buffer (Cat#9803S, Cell Signaling Technology) with protease inhibitor (Cat#11697698001, Sigma-Aldrich) and phosphatase inhibitor cocktails (Cat#524627-1EA, EMD Millipore) followed by standard Western blot assays. Primary antibodies include rabbit anti-phosphorylated STAT3 at Tyr705 (1:1000, Cat#9145S, Cell Signaling Technology), rabbit anti-STAT3 (1:1000, Cat#12640S, Cell Signaling Technology), rabbit anti-Cleaved Caspase-3 (Asp175) (1:1000, Cat#9661S, Cell Signaling Technology), and mouse anti-β-actin (1:2000, Cat#A5441, Sigma Aldrich). HRP-conjugated secondary antibodies include goat anti-rabbit (1:3000; Santa Cruz Biotechnology, sc-2004) and goat anti-mouse (1:3000; BD Biosciences, 554002). The antigen-antibody complex was detected by SuperSignal™ West Dura Extended Duration Chemiluminescent Substrate (Cat#34075, Thermo Fisher).

### Immunohistochemistry of human lung tissues

Paraffin sections (4 μm) of human lung samples were processed for immunostaining according to standard protocols. Primary antibodies include rabbit anti-RFX3 (1:100, Cat#HPA035689, Sigma Aldrich), rabbit anti-KRT5 (1:100, Cat# ab52635, Abcam), rabbit anti-Cleaved Caspase-3 (Asp175) (1:100, Cat# 9661, Cell Signaling Technology), and rabbit anti-SARS-CoV nucleocapsid protein (N) antibody that cross-reacts with SARS-CoV-2 N (1:2000, Cat# 200-401-A50, Rockland Immunochemicals). Biotinylated goat anti-rabbit (IgG, 1:200, Cat#BA-1000, Vector Laboratories) was used as the secondary antibody. For DAB staining, sections were treated with standard ABC kit (Vector Labs, PK-6100) and DAB Peroxidase Substrate Kit (Vector Labs, SK-4100). After DAB staining, samples were counterstained with Hematoxylin QS (Vector Laboratories, H-3404). Bright field images were taken using a digital camera (Nikon DS-Fi2).

## QUANTIFICATION AND STATISTICAL ANALYSIS

### Imaging and quantification

Stained slides were examined with a Nikon Ti inverted fluorescence/confocal microscope or Zeiss AX10 brightfield microscope. All images were processed using ImageJ software. Quantification was performed by two independent and blinded examiners as follows. For ALI and lung sections, 4-6 individual 20X images per sample were randomly selected and captured. The number of epithelial cells per image was counted based on the nuclei countered stained with DAPI or hematoxylin. Epithelial cells labelled with lineage-specific markers or RSV F protein in the same image were counted and reported as the percentage of the epithelial cell population of interest.

### Statistical analysis

For details on statistical analysis, the numbers of samples, and the experimental repeat, see corresponding figure legends and Results section. For statistical comparison between two experimental groups, unpaired Student’s t test was applied. For statistical comparison between more than 2 groups, one-way ANOVA followed Dunnett’s multiple comparison test or two-way ANOVA followed by Sidak’s multiple comparison test were statistically appropriate. These tests were performed using GraphPad Prism 8. A calculated p value less than 0.05 is considered statistically significant.

