## Supplemental Table and figures for "Age-related STAT3 signaling regulates severity of respiratory syncytial viral infection in human bronchial epithelial cells"

### SUPPLEMENTARY INFORMATION

**Supplementary Table 1. Demographic information of patients from whom TA BSCs were derived.**

| **Neonatal BSC lines** | | | | |
|---|---|---|---|---|
| Patient Study ID | Gestational Age (weeks) | Sex | Disease History | Assays performed |
| N1 | 39 | M | NAS | RS, DI, RI |
| N2 | 41 | F | HIE | RS, DI, RI, SI |
| N3 | 38 | F | Fetal maternal hemorrhage, polyhydramnios, PPHN | RS, DI |
| N4 | 41 | M | MAS, PPHN, HIE | RS, DI, RI |
| N5 | 40 | M | Pneumothorax | RS, DI, RI, SI |
| N6 | 41 | M | Amniotic fluid aspiration, PNA | RS, DI, RI |
| N7 | 39 | F | MAS, PPHN | RS, DI |
| N8 | 40 | M | MAS, PPHN, pulmonary hemorrhage | RS, DI |
| N9 | 39 | F | Feeding problem | DI, RI |
| N10 | 37 | M | Feeding problem, seizures | DI, RI |
| **Adult BSC lines** | | | | |
| Patient Study ID | Age (years) | Sex | Disease History | Assays performed |
| A1 | 36 | F | No known respiratory disease | RS, DI, RI, SI |
| A2 | 37 | M | Former smoker | RS, DI, RI |
| A3 | 32 | M | Former smoker | RS, DI |
| A4 | 45 | M | Former smoker | RS, DI, RI |
| A5 | 55 | F | No known respiratory disease | DI, RI, SI |

***NAS:** neonatal abstinence syndrome, **HIE:** hypoxic-ischemic encephalopathy, **PPHN:** persistent pulmonary hypertension of the newborn, **PNA:** pulmonary nodular amyloidosis, **MAS:** macrophage activation syndrome

***RS** = RNA Sequencing, **DI** = Differentiation, **RI** = RSV infection, **SI** = SARS-CoV-2 infection

**Supplementary Table 2.** List of primers used for RT-qPCR.

| Gene | Forward Primer | Reverse Primer |
|---|---|---|
| *RSV L2* | GAACTCAGTGTAGGTAGAATGTTTGCA | TTCAGCTATCATTTTCTCTGCCAAT |
| *CXCL10* | GAAATTATTCCTGCAAGCCAATTT | TCACCCTTCTTTTTCATTGTAGCA |
| *BCL2* | GGTGGGGTCATGTGTGTGG | CGGTTCAGGTACTCAGTCATCC |
| *MCL1* | GTGCCTTTGTGGCTAAACACT | AGTCCCGTTTTGTCCTTACGA |
| BIRC5 | AGGACCACCGCATCTCTACAT | AAGTCTGGCTCGTTCTCAGTG |
| *BCL2L1* | GAGCTGGTGGTTGACTTTCTC | TCCATCTCCGATTCAGTCCCT |
| *IFNL1* | AATTGGGACCTGAGGCTTCTC | CCAGCGGACTCCTTTTTGG |
| *IFNL3* | TAAGAGGGCCAAAGATGCCTT | CTGGTCCAAGACATCCCCC |
| *GAPDH* | GGAGCGAGATCCCTCCAAAAT | GGCTGTTGTCATACTTCTCATGG |
| *TNFA* | GAGGCCAAGCCCTGGTATG | CGGGCCGATTGATCTCAGC |
| *IFNB* | ATGACCAACAAGTGTCTCCTCC | GGAATCCAAGCAAGTTGTAGCTC |
| *MUC5AC* | CTCCTACCAATGCTCTGTA | GTTGCAGAAGCAGGTTTG |
| *MUC5B* | GACAGAGACGACAATGAG | CCTGATGTTTTCAAAAGTTTC |
| **shRNA target sequence** | | |
| *STAT3 shRNA* | GGCGTCCAGTTCACTACTAAA | |

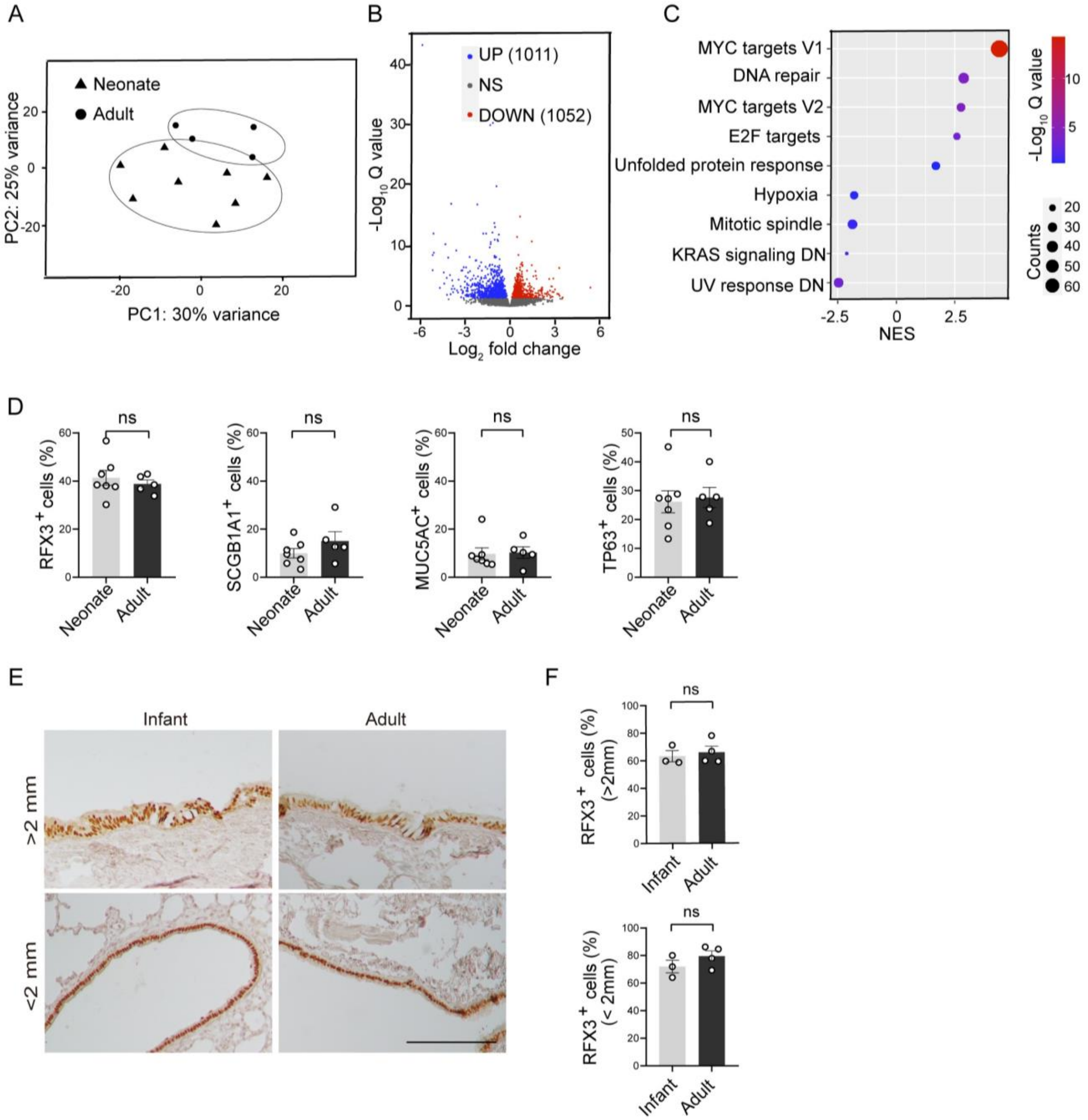


**Figure S1. BSCs from infants and adults have similar differentiation potentials. Related to Figure 1.**

(A) Principal component (PC) analysis of bulk RNA sequencing datasets of TA BSC lines from neonates (n=8) and adults (n=4). (B) Volcano plot showing the number of differentially expressed genes in neonatal BSCs compared to adult BSCs (padj<0.05). (C) Pathway analysis of bulk RNA sequencing results of neonatal and adult TA BSCs. (D) The relative abundance of different epithelial cell types based on antibody staining of day 21 ALI cultures of neonatal (n=7) and adult (n=5) TA BSCs. (E) Representative RFX3 staining of healthy human donor lungs from infants (n=3) and adults (n=4). Scale bar, 200 μm. (F) The relative abundance of $RFX3^+$ ciliated cells in human donor lungs of infants and adults. Each dot represents one BSC line in (D) or one donor in (F). Bar graphs show mean ± SEM. ns, not significant by Student's t-test (two-tailed) (D and F).

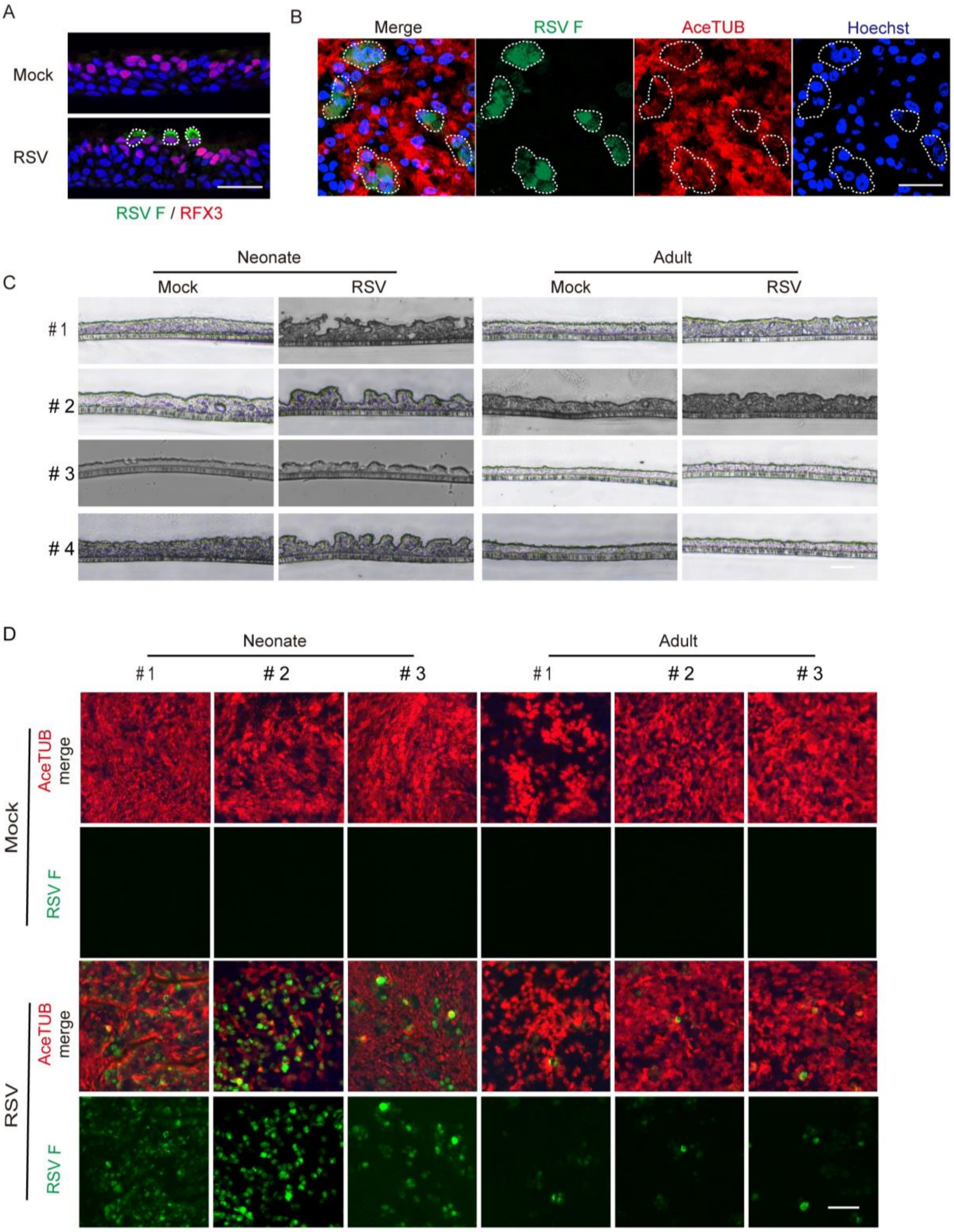


**Figure S2. Neonatal and adult ALI cultures show age-related severity of RSV infection. Related to Figure 1.**

(A) Representative double staining for RFX3 and RSV F in adult ALI cultures at 2 dpi. (B) Representative whole-mount staining for AceTUB and RSV F in adult ALI cultures at 2 dpi. Ciliated cells infected with RSV were outlined. (C) Representative brightfield images of neonatal (n=4) and adult (n=4) ALI cultures with and without RSV infection at 2 dpi. (D) Representative whole-mount staining for AceTUB and RSV F in neonatal (n=3) and adult (n=3) ALI cultures with and without RSV infection at 2 dpi. Scale bars, 50 μm.

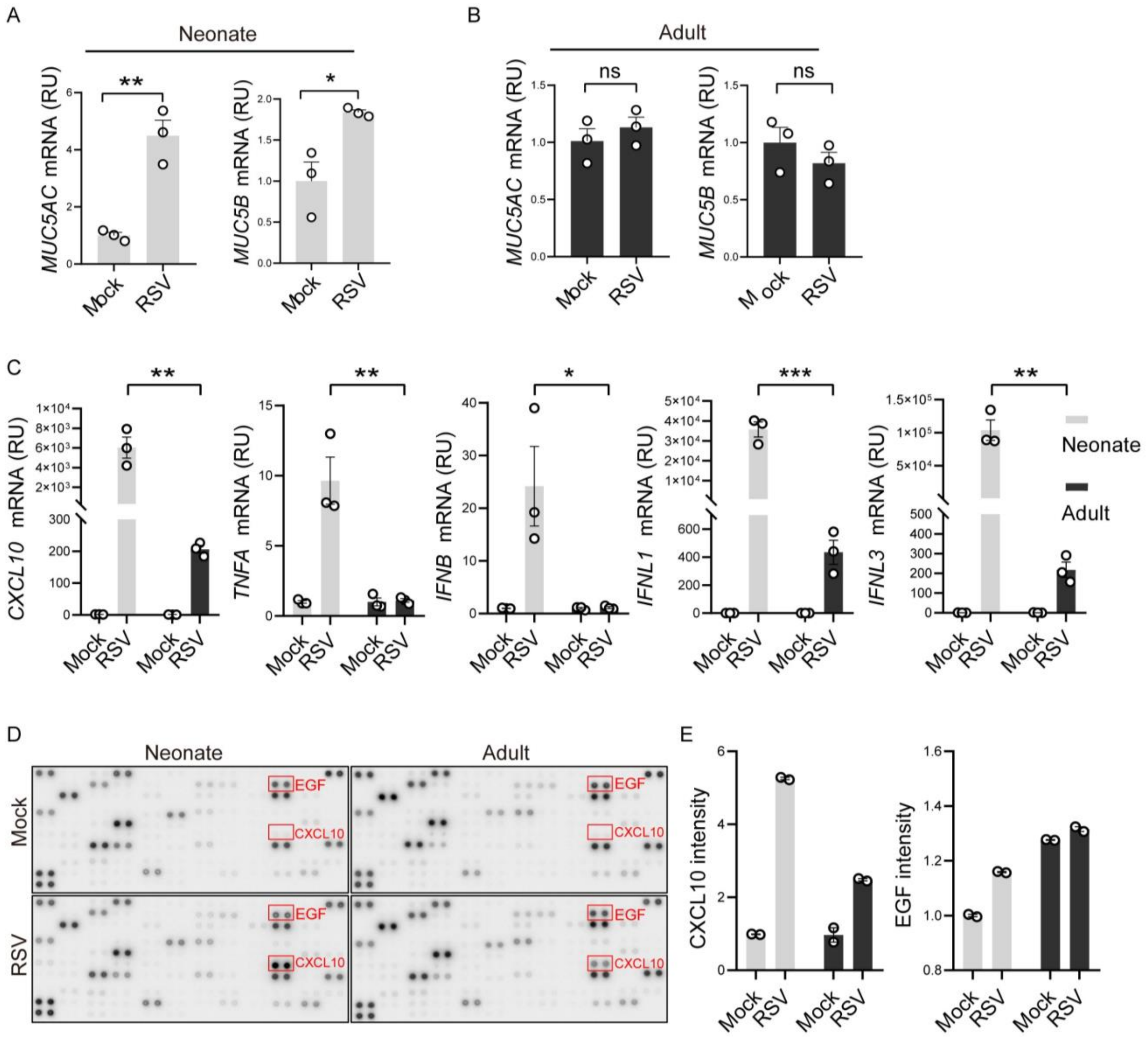


**Figure S3. RSV infection elevates more robustly mucin and cytokine/chemokine gene expression in neonatal ALI cultures than adult ALI cultures at 2 dpi. Related to Figure 1.**

(A-C) The relative levels of *MUC5AC*, *MUC5B, CXCL10*, *TNFA*, *IFNB*, *IFNL1*, and *IFNL3* gene expression in neonatal and adult ALI cultures at 2 dpi by RT-qPCR. (D) Human cytokine arrays using the medium collected from the bottom chamber in neonatal and adult ALI cultures at 2 dpi. (E) Densitometry measurements of the relative levels of CXCL10 and EGF. Each dot represents one BSC line. Bar graphs show mean ± SEM. *p<0.05, **p<0.01, and ***p<0.001 calculated by Student's t-test (two-tailed) in (A and B) and two-way ANOVA followed by Sidak's multiple comparison test in (C).

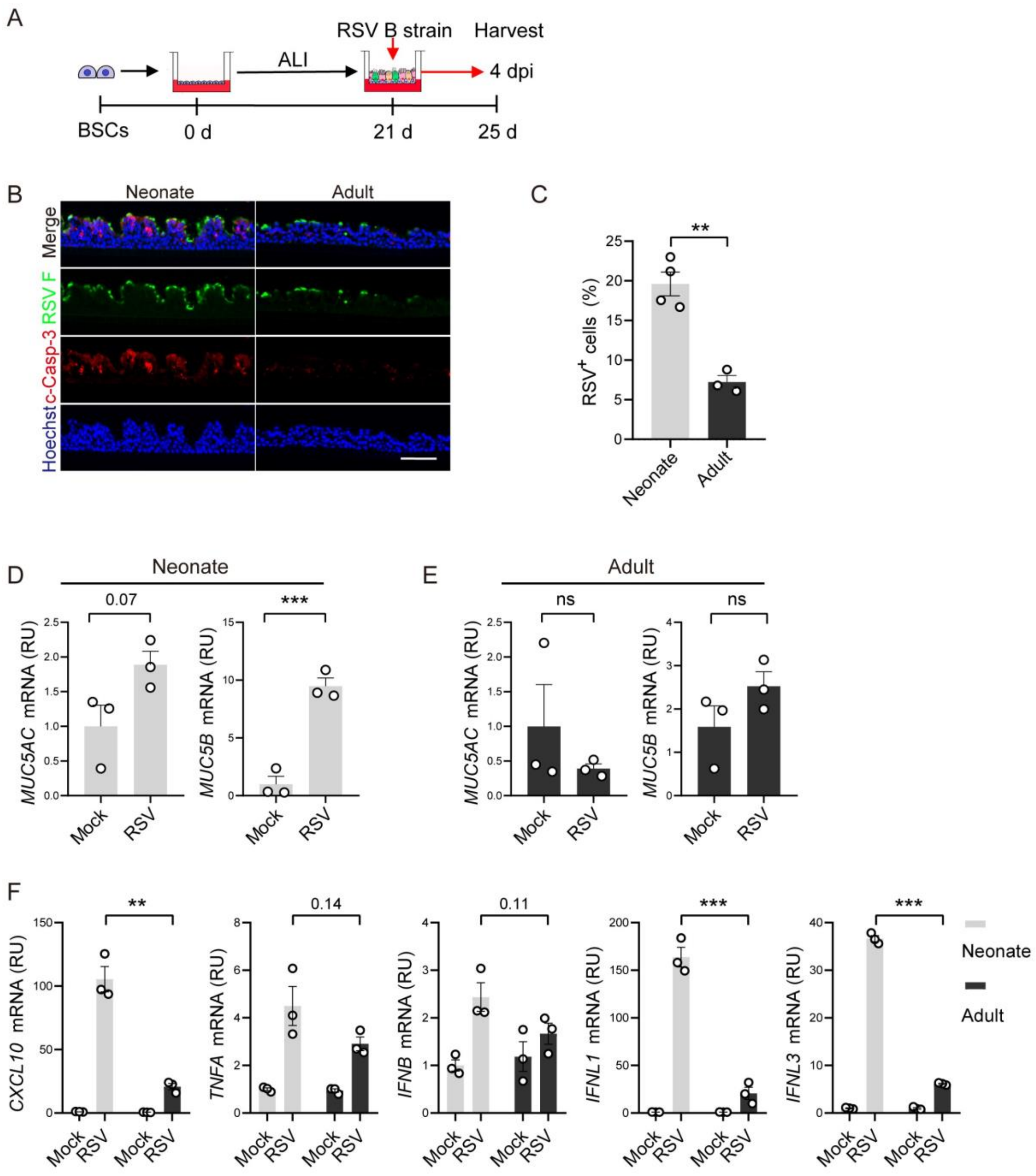


**Figure S4. RSV strain B induces age-related severity of infection in neonatal bronchial epithelium model. Related to Figure 2.**

(A) Schematic of infection of neonatal and adult ALI cultures with RSV B virus (WV/14617/85, MOI 2). Assays were performed at 4 dpi. (B) Representative double staining for RSV F protein and c-Casp-3. (C) The relative abundance of RSV $F^{+}$ cells. (D-F) The relative mRNA levels of *MUC5AC*, *MUC5B, CXCL10*, *TNFA*, *IFNB*, *IFNL1*, and *IFNL3* by RT-qPCR.

Each dot represents one BSC line. Bar graphs show mean ± SEM. *p<0.05, **p<0.01, and ***p<0.001 calculated by Student's t-test (two-tailed) in (C-E) and two-way ANOVA followed by Sidak's multiple comparison test in (F).

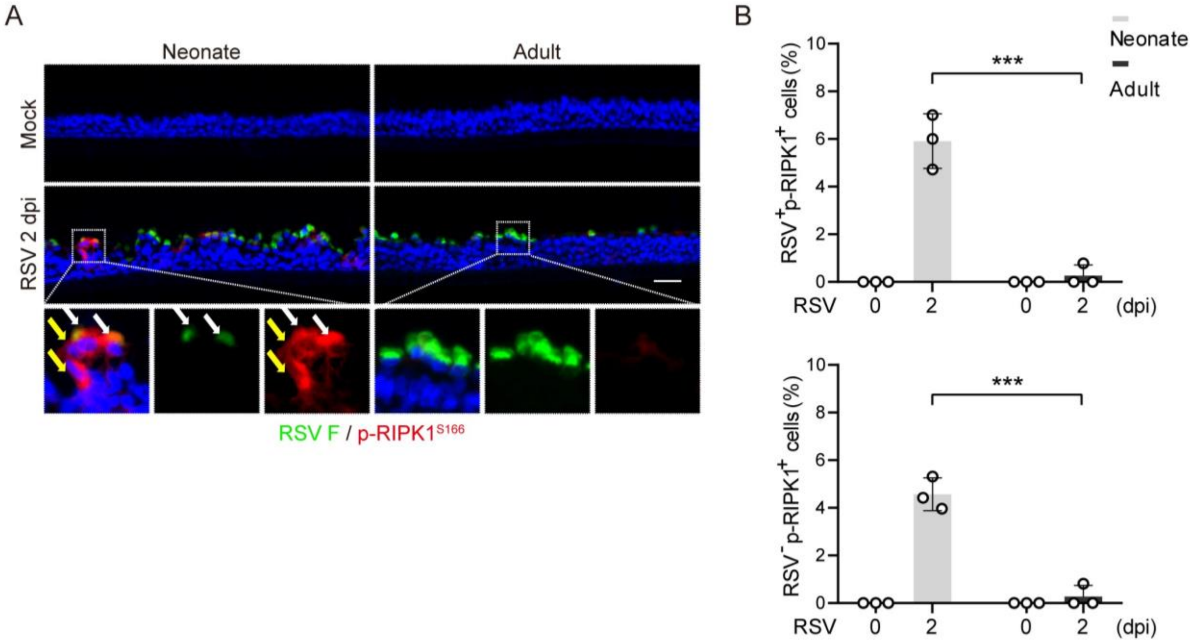


**Figure S5. RSV infection induces necroptosis in a small number of epithelial cells in neonatal ALI cultures. Related to Figure 2.**

(A) Representative double staining for RSV F and p-RIPK1$^{S166}$ using sections of neonatal and adult ALI cultures (n=3 BSC lines per age group) at 2 dpi. White arrows mark RSV F$^{+}$p-RIPK1$^{+}$ cells and yellow arrows mark RSV F$^{-}$p-RIPK1$^{+}$ cells.

(B) The relative abundance of RSV F$^{+}$ p-RIPK1$^{+}$ and RSV F$^{-}$ p-RIPK1$^{+}$ cells.

Each dot represents one BSC line. Bar graphs represent mean ± SEM. Statistical significance was calculated by two-way ANOVA followed by Sidak's multiple comparison test in (B). ***$p<0.001$. Scale bar, 50 μm.

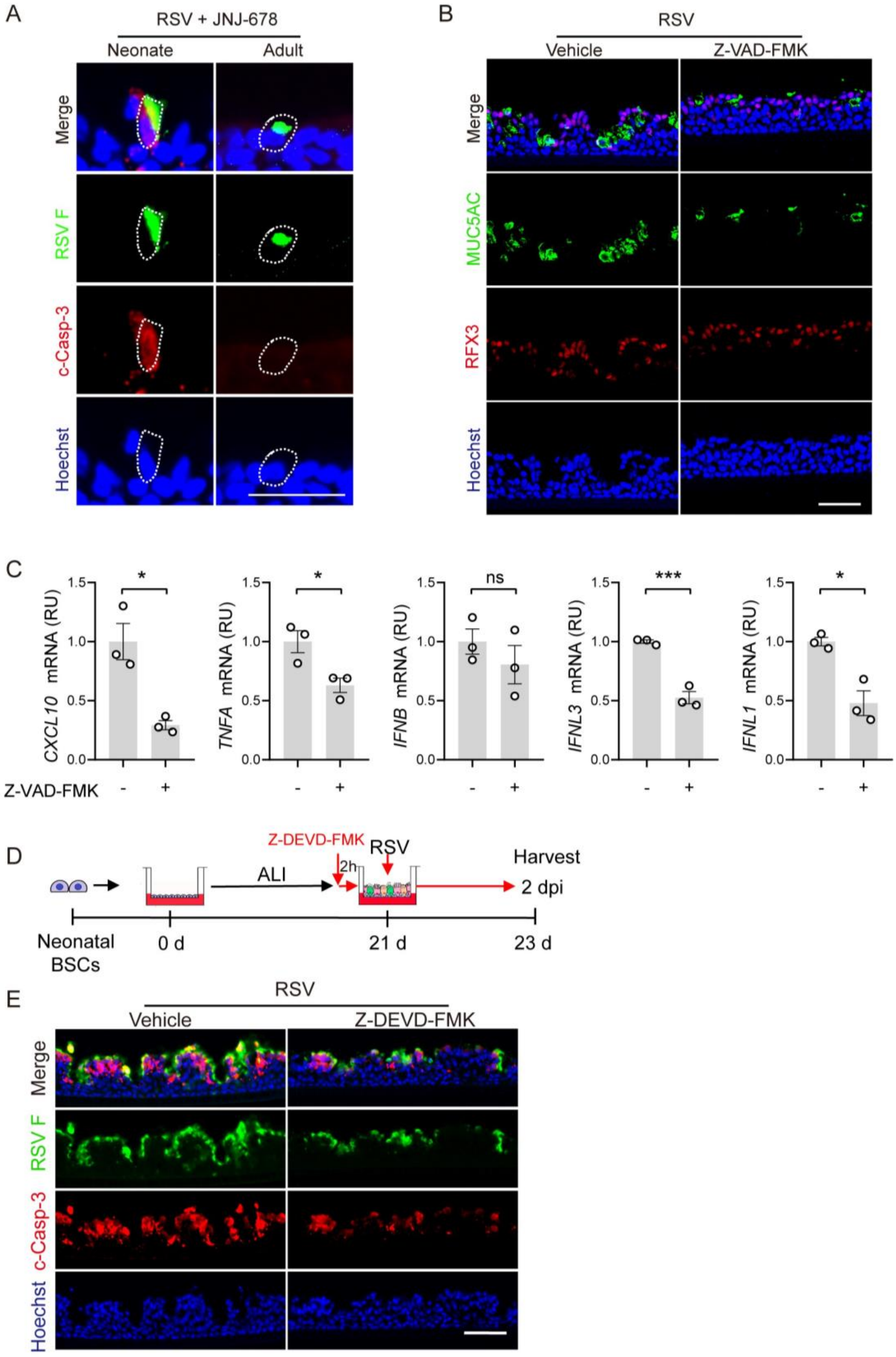


**Figure S6. Blockade of apoptosis in neonatal bronchial epithelium following RSV infection reduces mucus hyperplasia and cytokine/chemokine gene expression. Related to Figure 2.**

(A) Representative double staining for RSV F and c-Casp-3 in JNJ-678-treated neonatal ALI cultures at 2 dpi. JNJ-678 treatment was described in Figure 1J. (B) Representative double staining for MUC5AC and RFX3 neonatal ALI cultures treated with Z-VAD-FMK at 2 dpi. Z-VAD-FMK treatment was described in Figure 2E. (C) The relative levels of *CXCL10*, *TNFA*, *IFNB*, *IFNL1*, and *IFNL3* expression with and without Z-VAD-FMK treatment at 2 dpi by RT-q-PCR. (D) Schematic of Z-DEVD-FMK (40 μM) treatment. (E) Representative double staining for RSV F and c-Casp-3 with and without Z-DEVD-FMK treatment at 2 dpi.

Each dot represents one BSC line. Bar graphs show mean ± SEM. *p<0.05 and ***p<0.001 calculated by Student's t-test (two-tailed). Scale bars, 50 μm.

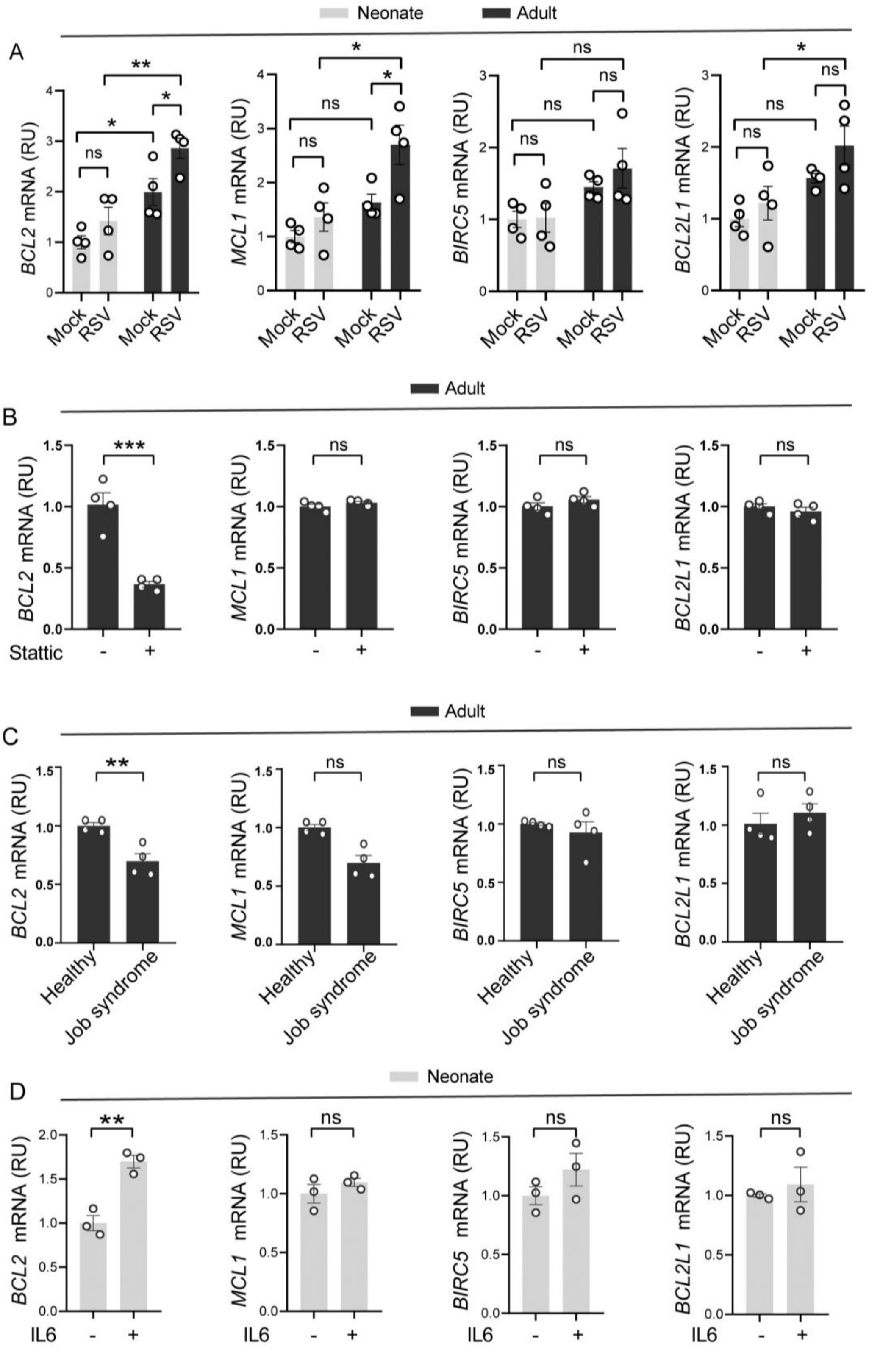


**Figure S7. *BCL2* family gene expression associated with RSV infection and STAT3 signaling in neonatal and adult ALI cultures. Related to Figures 3, 6, and 7.**

(A-D) The relative levels of *BCL2*, *MCL1*, *BIRC5*, and *BCL2L1* expression in neonatal and adult ALI cultures with and without RSV infection in (A), Stattic (20 μM)-treated adult ALI cultures in (B), adult ALI cultures of BSCs from healthy donors and a patient with Job syndrome harboring STAT3-S560del mutation (C), neonatal ALI cultures with and without IL6 (50 ng/mL) treatment in (D).

Each dot represents one BSC line in (A) or one of 4 independent experiments of one BSC line in (B, C, and D). Bar graphs show mean ± SEM. *p<0.05, **p<0.01, and ***p<0.001 calculated by two-way ANOVA followed by Sidak's multiple comparison test in (A) and by Student's t-test (two-tailed) in (B-D).

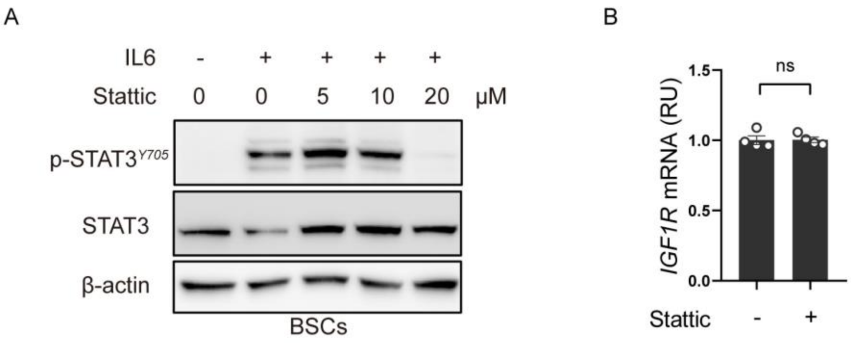


**Figure S8. Dose-dependent blockade of IL6-induced STAT3 activation by Stattic. Related to Figure 6 and 7.**

(A) Representative Western blot assay for the level of p-STAT3$^{Y705}$ and STAT3 in BSCs pretreated with Stattic at different concentrations 2 h prior to IL-6 stimulation (50 ng/mL). Samples were collected 30 min after IL6 stimulation. β-Actin was loading control. (B) The relative level of *IGF1R* gene expression in adult ALI cultures with and without Stattic treatment for 48 hours by RT-qPCR. Stattic (20 μM) was given to the bottom chamber.

Each dot represents one of 4 independent experiments of one BSC line in (B). Bar graphs show mean ± SEM. ns, not significant by Student’s t-test (two-tailed).

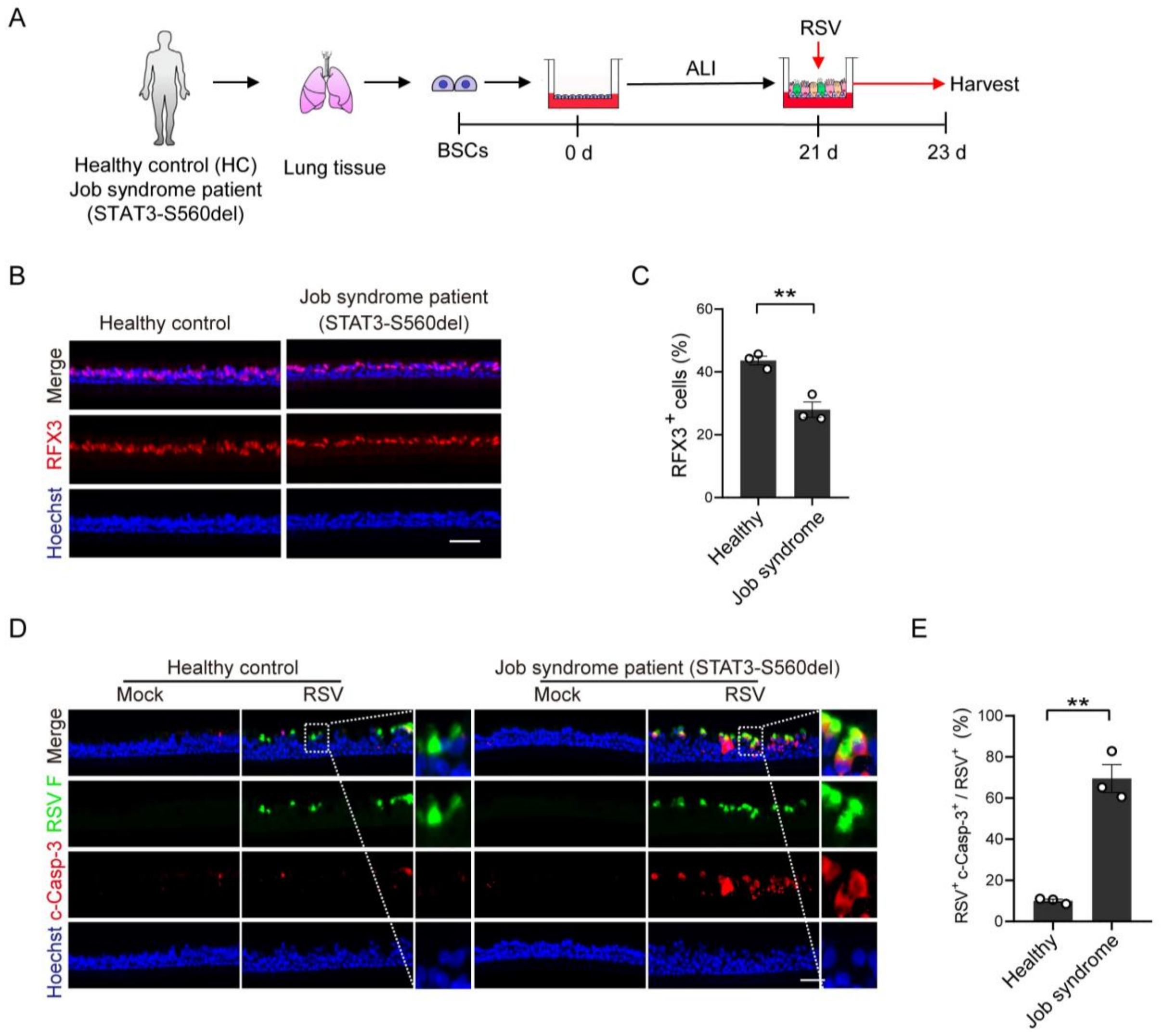


**Figure S9. The ALI culture derived from BSCs from an adult patient with Job syndrome shows worsened apoptosis following RSV infection. Related to Figure 6.**

(A) Schematic of ALI cultures generated with BSCs isolated from lung biopsy samples of healthy donors and a patient with Job syndrome harboring STAT3-S560del mutation. (B) Representative RFX3 staining in healthy and Job syndrome ALI cultures. (C) The relative abundance of RFX3$^{+}$ ciliated in healthy and Job syndrome ALI cultures. (D) Representative double staining for RSV F and c-Casp-3 in healthy and Job syndrome ALI cultures at 2 dpi. (E) The relative abundance of RSV F$^{+}$c-Casp-3$^{+}$ double positive cells in RSV$^{+}$ cells in these cultures.

Data points represent 3 independent experiments of one BSC line for each group in (C and E). Bar graphs show mean ± SEM. **p<0.01 and ***p<0.001 calculated by Student’s t-test (two-tailed). Scale bars, 50 µm.

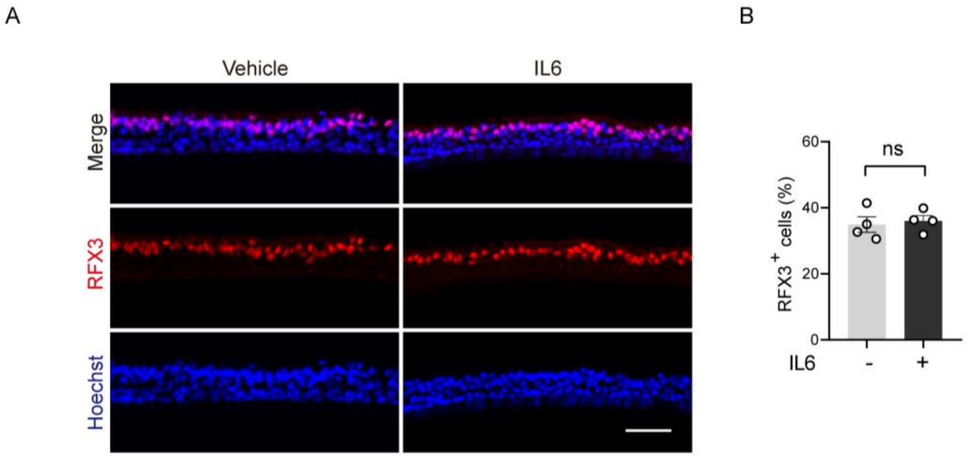


**Figure S10. IL6 treatment starting from day 18 in ALI has no effect on ciliated cell differentiation. Related to Figure 7.**

(A) Representative RFX3$^+$ staining in ALI cultures treated with vehicle or IL6 (50 ng/mL) starting from day 18 in ALI. ALI cultures were assayed at day 21. (B) The relative abundance of RFX3$^+$ ciliated with and without IL6 treatment.

Data points represent 4 independent experiments of one neonatal BSC line. Bar graph shows mean ± SEM. ns, not significant calculated by Student's t-test (two-tailed). Scale bar, 50 μm.
